# Periductal Fibroblast Density Defines Lymphocyte Exclusion via a CD44-Dependent Stromal Checkpoint in Pancreatic Cancer

**DOI:** 10.1101/2024.05.23.593868

**Authors:** Zoe X. Malchiodi, Alexander A. Lekan, Robert K. Suter, Atul Deshpande, Ivana Peran, Brent T. Harris, Anju Duttargi, Min-Ju Chien, Samika Hariharan, Lucia Wetherill, Selime Arslan, Sandra A. Jablonski, Won Jin Ho, Marwa Afifi, Elana J. Fertig, Louis M. Weiner

**Author notes:** These authors contributed equally to this work (Zoe X. Malchiodi, Alexander A. Lekan). **Address correspondence to:** Louis M. Weiner, MD, Georgetown Lombardi Comprehensive Cancer Center, Research Building E501, 3970 Reservoir Rd NW, Washington, DC, 20057.

## Abstract

Pancreatic ductal adenocarcinoma (PDAC) exhibits dense fibrosis and immune exclusion. While fibrosis has been studied globally and at the region-of-interest level, its impact on stromal-ductal architecture and immune cell localization remains unknown. Here, we establish cancer-associated fibroblast (CAF)-stratified ductal spatial architecture as a fundamental determinant of immune exclusion in PDAC. Focusing on malignant PDAC epithelial ductal regions, the critical interface where immune cells must access tumor epithelium, we demonstrate that periductal fibroblast organization dictates leukocyte proximity. Through integrative analysis of treatment-naïve patient samples from three independent cohorts – including imaging mass cytometry, multiplex immunohistochemistry, and single-cell RNA sequencing – we uncovered that activated, pro-inflammatory leukocytes preferentially localized near malignant ducts in regions with low fibroblast density. Stratifying epithelial-ductal regions by CAF abundance revealed a graded constraint: increasing fibroblast content corresponded to reduced leukocyte–epithelial proximity and elevated collagen I deposition. Despite their exclusion in high-CAF ducts, leukocytes in low-CAF ducts retained functional competence. Mechanistically, ligand–receptor inference implicated collagen–CD44 signaling as an adhesion axis anchoring immune cells within fibroblast-rich zones, with CD44 blockade enhancing natural killer cell invasion and motility in fibrotic spheroid models. Thus, by establishing ductal regions as critical spatial units of immune exclusion, these findings provide a framework for dissecting stromal–immune interactions and reveal targetable “stromal checkpoints” that can be leveraged to overcome CAF-driven barriers to leukocyte motility and infiltration in PDAC.

**Graphical Abstract:** 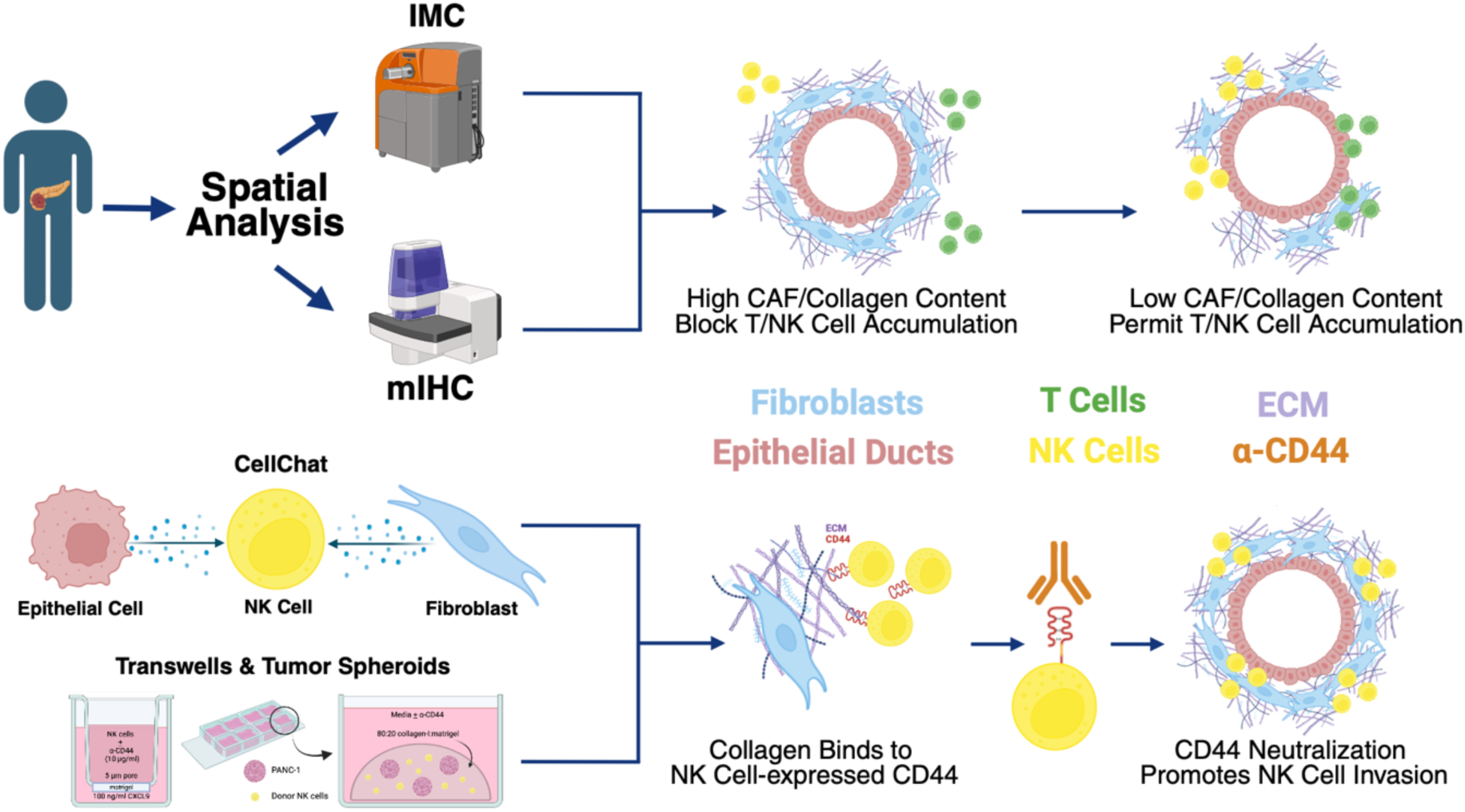

## Introduction

Pancreatic ductal adenocarcinoma (PDAC) has a 5-year survival rate of 12%^1^. PDAC is considered to be an immunologically “cold” tumor with minimal infiltration by activated leukocytes and immunotherapy-based clinical trials using checkpoint inhibitors have been largely unsuccessful^2–4^. This resistance can be at least partly attributed to the tumor’s desmoplastic stroma, driven by extracellular matrix (ECM) deposition from cancer-associated fibroblasts (CAFs)^5^. The resulting fibrotic barrier increases tissue stiffness while restricting activated leukocyte infiltration and access to malignant ductal epithelial cells^6, 7^. Therefore, defining how CAF spatial organization within the periductal stroma constrains activated leukocyte-epithelial cell interactions is critical to creating successful immune-based therapeutic approaches.

Recent studies have explored the impact of CAFs on tumor cell progression in PDAC and characterized the landscape of the PDAC TME using multiple spatial molecular profiling technologies^8–18^. While these studies have provided insights into immune dysfunction, CAF subtype heterogeneity, and desmoplastic remodeling, they have primarily focused on global tissue-level patterns or pre-selected regions (i.e. juxtatumoral vs. panstromal). Furthermore, the complex cellular interactions between CAFs, immune cells, and tumor cells within PDAC malignant ductal regions remain unknown. These ducts are the sites of tumor origin, progression, and metastasis that shape the global PDAC TME and its consequent biology, making the study of cellular colocalization and signaling in these sites critical to understand PDAC carcinogenesis^19–22^. Since these ductal regions represent the critical interface between malignant epithelium and infiltrating immune cells, defining the stromal barriers that restrict immune access in these regions is crucial for understanding and overcoming immune exclusion in PDAC.

Here, we identified periductal CAF density in malignant ducts within invasive PDAC as a key tumor organizational feature that dictates immune access to tumor epithelium. Using complementary spatial approaches, including imaging mass cytometry (IMC) and multiplex immunohistochemistry (mIHC), in separate cohorts of PDAC samples, we mapped periductal cellular interactions and found that periductal fibroblast organization is the principal determinant of leukocyte infiltration in malignant ductal regions. Increased periductal fibrosis correlated with increased collagen I deposition and increased proliferation of epithelial cells and corresponded with reduced leukocyte-epithelial proximity, particularly with regards to natural killer (NK) cells. Ligand-receptor inference analysis of a recently published scRNA-seq dataset^23^ of PDAC samples from 16 patients revealed increased collagen signaling from CAFs to leukocytes via CD44, and CD44 neutralization *in vitro* increased NK cell motility through the ECM. These results indicate that (1) periductal CAF architecture governs leukocyte access while promoting malignant epithelial cell proliferation and (2) targeting stromal checkpoints, such as CD44, may improve leukocyte infiltration and overcome ECM-mediated immune exclusion. Collectively, CAF-stratified ductal spatial architecture defines a new framework for analyzing PDAC fibrotic restrictions to leukocyte infiltration that can potentially be overcome through neutralization of stromal checkpoints in PDAC and other highly fibrotic tumors.

## Results

### Fibroblast composition organizes spatial architectures that determine immune access to malignant epithelial ducts in PDAC

We first performed IMC on a tissue microarray (TMA) of primary tumor samples from 21 treatment-naïve PDAC patients to define the major cellular compartments of the tumor microenvironment (TME). IMC is a high-dimensional spatial proteomics platform that maintains tissue architecture while capturing single-cell protein levels^24^. We used a custom 40-marker metal-conjugated antibody panel targeting cells from epithelial, immune and stromal lineages (**Supplementary Table S1).** Following imaging and cell segmentation, we identified 412,862 cells in 44 IMC pseudoimages, with two or more primary tumor cores being imaged for some patient **(Supplementary Table S2)**. Using a Seurat-based analysis pipeline^25–29^, single-cell protein expression data were log-normalized, and cell populations were identified using clustering approaches^30^, which resolved eight principal cell types encompassing malignant epithelial cells, cancer-associated fibroblasts (CAFs), leukocytes, and other stromal populations (**Supplementary Fig. S1**). Subclustering distinguished functional states: active, proliferative, or inhibitory, within each lineage (**Supplementary Fig. S2**). This cellular map established the framework for subsequent analyses of how CAF organization around malignant ducts constrains leukocyte access within the PDAC TME.

Having defined major cellular lineages by IMC, we next examined how these populations are spatially organized within the PDAC TME. Using cellular neighborhood analysis^31^, we classified distinct multi-cellular structures based on their cellular composition and proximity^31^. This analysis identified eight distinct cellular neighborhoods (**Figs. 1A-B**) each defined by characteristic combinations of epithelial, fibroblast, and immune populations. Fibroblast composition influenced neighborhood architecture: FAP^+^ fibroblast-rich regions (neighborhood 6) were associated with proliferative epithelial areas, while αSMA^+^ fibroblast-rich regions (neighborhood 7) were enriched for CD4^+^/CD8^+^ T cells and macrophages (**Fig. 1C**). In contrast, immune-enriched neighborhood 4 showed the majority of apoptotic epithelial cells and was largely depleted of FAP^+^ or αSMA^+^ fibroblasts, consistent with reduced stromal barriers in immune-accessible regions (**Fig. 1C, Supplementary Fig. S3)**. Notably, neighborhood 2 was enriched for IFNγ^+^ positive epithelial cells, a marker of immune activation^32^ (**Fig. 1C, Supplementary Fig. S3**). These epithelial neighborhoods also contained a higher density of leukocytes relative to fibroblasts (**Supplementary Fig. S3)**, supporting that immune cell accumulation occurs preferentially in epithelial compartments with reduced stromal architecture. Consistent with previous reports in untreated PDAC^11^, adaptive lymphoid cells (CD4^+^/CD8^+^ T cells, B cells) were confined to their own distinct neighborhood (3) and were spatially segregated from epithelial rich regions (**Fig. 1C**). In contrast, activated NK cells, based on Granzyme B expression, were observed in epithelial-dominant neighborhoods (1, 2, 6, and 8), suggesting that innate immune cells mediate the primary immune interactions at malignant ducts.

**Figure 1:**
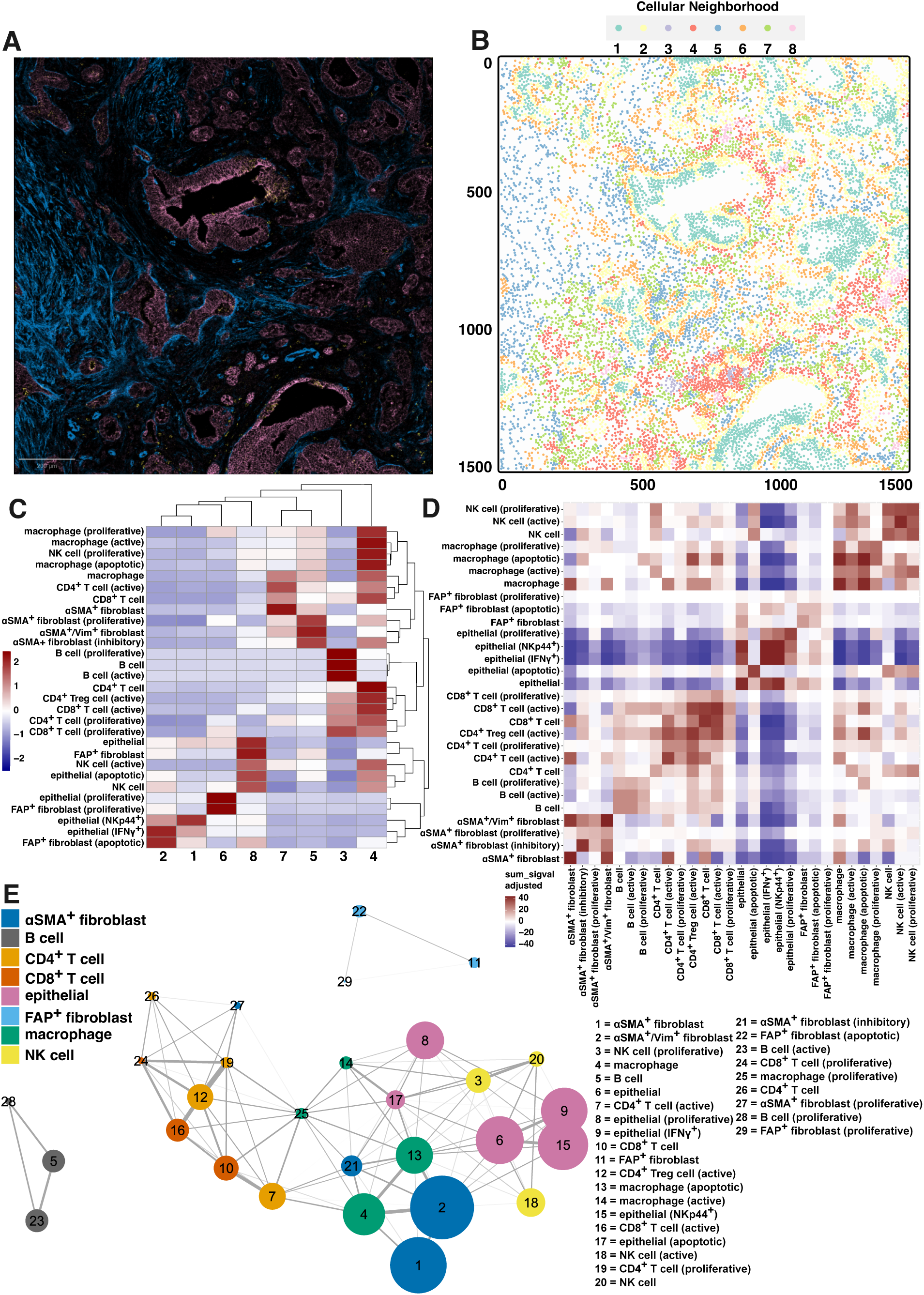
Fibroblast composition organizes spatial architectures that determine immune access to malignant epithelial ducts in PDAC. **A.** Corresponding patient IMC pseudoimage for CN spatial plot in **B** (αSMA = blue; E-cadherin, Pan-cytokeratin = pink; NKG2D = yellow, scale bar = 200 µm). **B.** Representative spatial plot of CNs in the human PDAC TME (CN1 = teal, CN2 = yellow, CN3 = purple, CN4 = red, CN5 = blue, CN6 = orange, CN7 = lime, CN8 = pink). **C.** Cellular neighborhood (CN) analysis revealed 8 distinct cellular neighborhoods (y-axis: cellular neighborhoods; red = high proportion of cell subpopulation of interest (x-axis) within cellular neighborhoods, blue = low proportion of cell subpopulation of interest within cellular neighborhoods). **C. D.** Cellular interaction analysis (blue = strong avoidance, red = strong interaction; FDR, *p* < 0.01) of IMC-defined cell subpopulations. **E.** Network graph of the top 75% of interactions between all IMC-defined cell populations in human PDAC showing epithelial cells and NK cells exist in similar networks.

Although CAF-mediated immune exclusion has been reported in PDAC^9, 16^, our analysis defines how distinct fibroblast subsets establish reproducible spatial architectures that dictate the degree of immune access to malignant ducts. Epithelial dominant neighborhoods (1, 2, 6, and 8) displayed variable leukocyte abundance depending on surrounding fibroblast density (**Fig. 1C**). Collectively, these analyses reveal that immune accessibility in PDAC is determined not by total stromal content, but by the local fibroblast organization encasing malignant epithelial ducts, prompting a focused analysis of epithelial-ductal regions to quantify how fibroblast abundance constrains leukocyte access within the TME.

Given that neighborhood analysis captures higher-order organization but not direct cellular interactions, we next applied a permutation-based interaction analysis^28^, which identifies significant deviations from a random cellular arrangement, to infer colocalization between PDAC cellular subsets identified via IMC. Specifically, this analysis reveals which cellular subsets are spatially interacting more or less frequently than expected due to random variation. As expected, epithelial cells, fibroblasts, and immune cells tended to localize with cells of the same type (**Fig. 1D**). Confirming neighborhood analysis, activated NK cells, but not T cells, frequently interacted with apoptotic epithelial cells (**Fig. 1D)** while activated CD4^+^ and CD8^+^ T cells primarily associated with one another rather than with epithelial or stromal cells (**Fig. 1D**). Macrophages, on the other hand, most often interacted with αSMA^+^ fibroblasts (**Fig. 1D**).

To visualize these multicellular relationships within the TME, we performed a cell–cell network analysis defined from the average distance of spatial neighbors^33^, which shows that macrophage and αSMA⁺ fibroblast nodes have the closest cross-lineage connectivity, whereas NK cell nodes clustered near epithelial cell nodes, distinct from CD4⁺/CD8⁺ T cell and B cell networks (**Fig. 1E**). This network structure suggest that fibroblast organization not only structures epithelial–stromal interfaces but also constrains immune cell localization within the PDAC TME.

Validation in an independent cohort of PDAC tumors (*n* = 17) using multiplex immunohistochemistry (mIHC) confirmed the spatial relationships observed by IMC (**Supplementary Fig. S4A**). Fibroblasts and epithelial cells constituted the majority of the PDAC TME (**Supplementary Fig. S4B**). Notably, αSMA^+^ fibroblast content was inversely correlated with NK cell, but not macrophage, abundance (**Supplementary Fig. S4C**). CD16^+^ NK cells and macrophages, consistent with a pro-inflammatory phenotype^34, 35^, were reduced with higher αSMA^+^ fibroblast content in tumors, supporting cell-cell network analysis and suggesting a selective exclusion of effector leukocytes in fibroblast-rich regions (**Fig. 1E**). Overall, these analyses demonstrate that fibroblast organization determines immune architecture in PDAC, limiting the spatial access of pro-inflammatory, cytotoxic leukocytes to malignant epithelial ducts.

### CAF-stratified ductal spatial architecture governs immune access in PDAC

Given that increased αSMA^+^ fibroblasts are inversely correlated with effector leukocyte infiltration (**Supplementary Fig. S4C**), we next sought to understand how per-ductal fibroblasts influence leukocyte infiltration in regions near the duct. Using epithelial-ductal regions of interest (ROIs) defined from IMC-based cellular neighborhood analysis, we applied Seurat’s^27^ CellSelector function to isolate and quantify immune and stromal populations across 952 epithelial-ductal ROIs. These ROIs were stratified into 4 quartiles (Q1-Q4) according to fibroblast abundance, with Q1 representing fibroblast-low and Q4 representing fibroblast-high ducts. This stratification revealed a consistent CAF-stratified ductal spatial architecture (**Figs. 2A-B**) where increasing fibroblast content corresponded to progressive restriction of leukocyte–epithelial proximity. Fibroblast abundance increased significantly across quartiles (**Fig. 2C**), driven predominantly by αSMA^+^ fibroblasts, rather than FAP^+^ fibroblasts (**Supplementary Fig. S5**), supporting previous reports that αSMA^+^ fibroblasts cluster in periglandular regions^36^. As expected, increasing fibroblast abundance was associated with a corresponding rise in Collagen I expression (**Fig. 2D**), consistent with fibroblast-driven matrix deposition^37^. While total epithelial cells decreased with increasing fibroblast content (**Fig. 2E**), proliferative epithelial cells increased (**Fig. 2E**), supporting the interpretation that periductal fibroblast infiltration supports epithelial cell growth.

**Figure 2:**
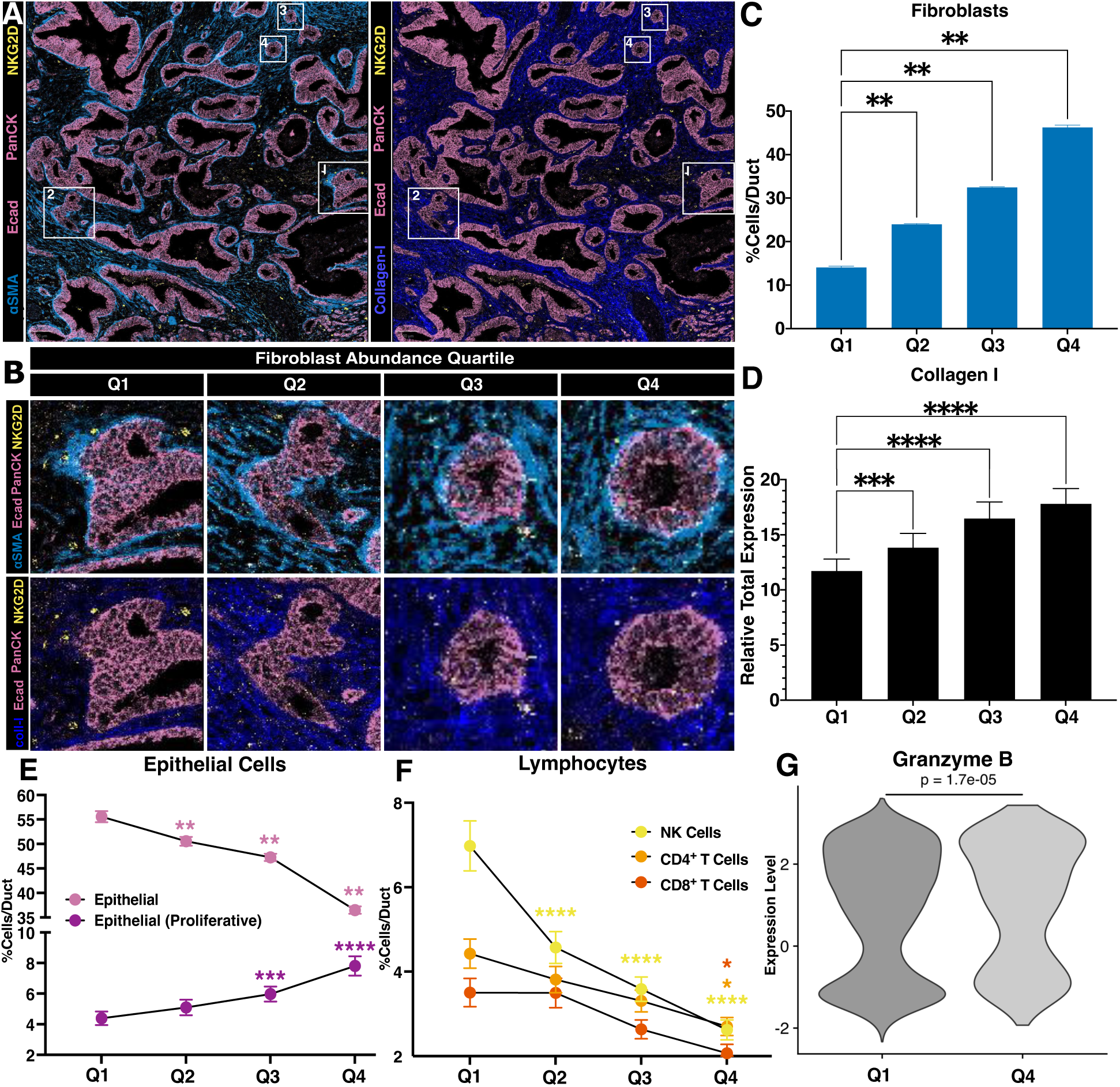
CAF-stratified ductal spatial architecture govern immune access in PDAC. **A.** Representative IMC pseudoimages showing αSMA (left, light blue) or collagen-I (right, dark blue) with pan-cytokeratin (panCK, pink), E-cadherin (Ecad, pink), and NKG2D (yellow); scale bar = 100 µm with white boxes of each fibroblast abundance quartile with corresponding numbers. **B.** Insets of representative epithelial-ductal ROIs from each fibroblast abundance quartile (Q1=238, Q2= 235, Q3=240, Q4=239) shown in the white insets in **A**. **C.** Bar plot of percent cells/ducts (mean ± SEM) of total fibroblasts, **D.** Bar plot of percent cells/ducts (mean ± SEM) of total collagen-I expression in epithelial-ductal ROIs ranked by fibroblast abundance quartile (2-way ANOVA; \**p* < 0.05, ** *p* < 0.01, \*\*\**p* < 0.001, \*\*\*\**p* < 0.0001; Q1=55, Q2= 58, Q3=57, Q4=60). **E**. Connecting line plot of percent cells/ducts (mean ± SEM) of total epithelial cells and proliferative epithelial cells (2-way ANOVA; \**p* < 0.05, \*\*\*\**p* < 0.0001; all quartiles compared to Q1). **F.** Connecting line plot of percent cells/ducts (mean ± SEM) of NK cells, CD4^+^ T Cells, and CD8^+^ T Cells (2-way ANOVA; \**p* < 0.05, \*\*\*\**p* < 0.0001; all quartiles compared to Q1). **G.** Violin plot showing NK cell-based expression in Q1 and Q4 fibroblast abundance quartiles of Granzyme B (Kruskal-Wallis test).

We next assessed the impact of fibroblasts on the spatial associations of activated leukocytes with ductal epithelial cells. Expression of NKG2D, an activating receptor expressed on human lymphocytes^38^, declined as fibroblast and collagen content increased (**Fig. 2B**). Among immune subsets, NK cells showed the strongest sensitivity to fibroblast density; their abundance was highest in fibroblast-low ducts (Q1) and decreased progressively across quartiles (**Fig. 2F**), indicating fibroblast-dependent exclusion of cytotoxic innate lymphocytes. In contrast, CD4⁺ and CD8⁺ T cell levels remained relatively stable (**Fig. 2F**), consistent with their preferential localization to stromal rather than malignant ductal epithelium^39^. The loss of NK cell infiltration was accompanied by reduced IFNγ⁺ epithelial cells (**Supplementary Fig. S6E**), implicating diminished NK-epithelial cell interactions and associated cytolytic signaling. In fibroblasts-sparse ducts(Q1-Q2), NK cells retained high expression of activation markers (CD69, CD107a, IFNγ) (**Supplementary Fig. S7**), and displayed significant increases in Granzyme B expression (**Fig. 2G**), confirming preserved effector function where malignant epithelial cell access is maintained.

Collectively, these findings establish CAF-stratified ductal spatial architecture as a defining architecture feature of PDAC, whereby periductal fibroblast density promotes malignant epithelial proliferation while restricting NK cell infiltration and activation. This represents a novel mechanism of immune evasion in PDAC and suggests that periductal stromal remodeling to reduce fibroblast density or collagen deposition may be required to enable effective lymphocyte-based immunotherapies.

### Cell-cell communication analysis implicates CD44 as a stromal checkpoint mediating NK-ECM interactions

Because NK cell access to malignant ducts was inversely correlated with periductal fibroblast density (**Fig. 2F–G**), we next sought to identify the molecular pathways that could mediate these fibroblast-NK cell interactions within the PDAC tumor microenvironment. We applied ligand-receptor inference of cell to cell communication with CellChat^40^ to a reference immune-enriched human PDAC scRNAseq dataset containing matched adjacent-normal (AdjNorm) and PDAC tumor samples^23^. To distinguish malignant from normal epithelial cells, we used CytoTRACE to infer differentiation states^41,42^. We also subclustered fibroblasts to resolve canonical CAF subsets, including myofibroblasts (myCAFs; *FAP, MMP11, HOPX, POSTN, COL12A1*)^9^, inflammatory (iCAFs; *CFD, DPT, AGTR1, CXCL12, CCL2*)^9^, antigen-presenting (apCAFs; *HLA-DRA, HLA-DPA1, HLA-DQA1*)^9^, normal fibroblasts, and stellate-like fibroblasts (*MYH11, RGS5)*^43, 44^ (**Supplementary Figs. S8A-E**).

Because NK cells were the immune population most affected by periductal fibroblast density (**Fig. 2G–H**), we next examined whether fibroblasts engage in molecular signaling that could underlie this selective exclusion. CellChat analysis revealed that CAFs in PDAC, particularly iCAFs and myCAFs, exhibited increased outgoing signaling compared with adjacent-normal fibroblasts, whereas NK cells and malignant epithelial cells served as prominent signal receivers (**Fig. 3A-B**). This suggests that CAF-derived signaling networks may directly modulate NK cell behavior and epithelial interactions, extending their role beyond structural confinement.

**Figure 3:**
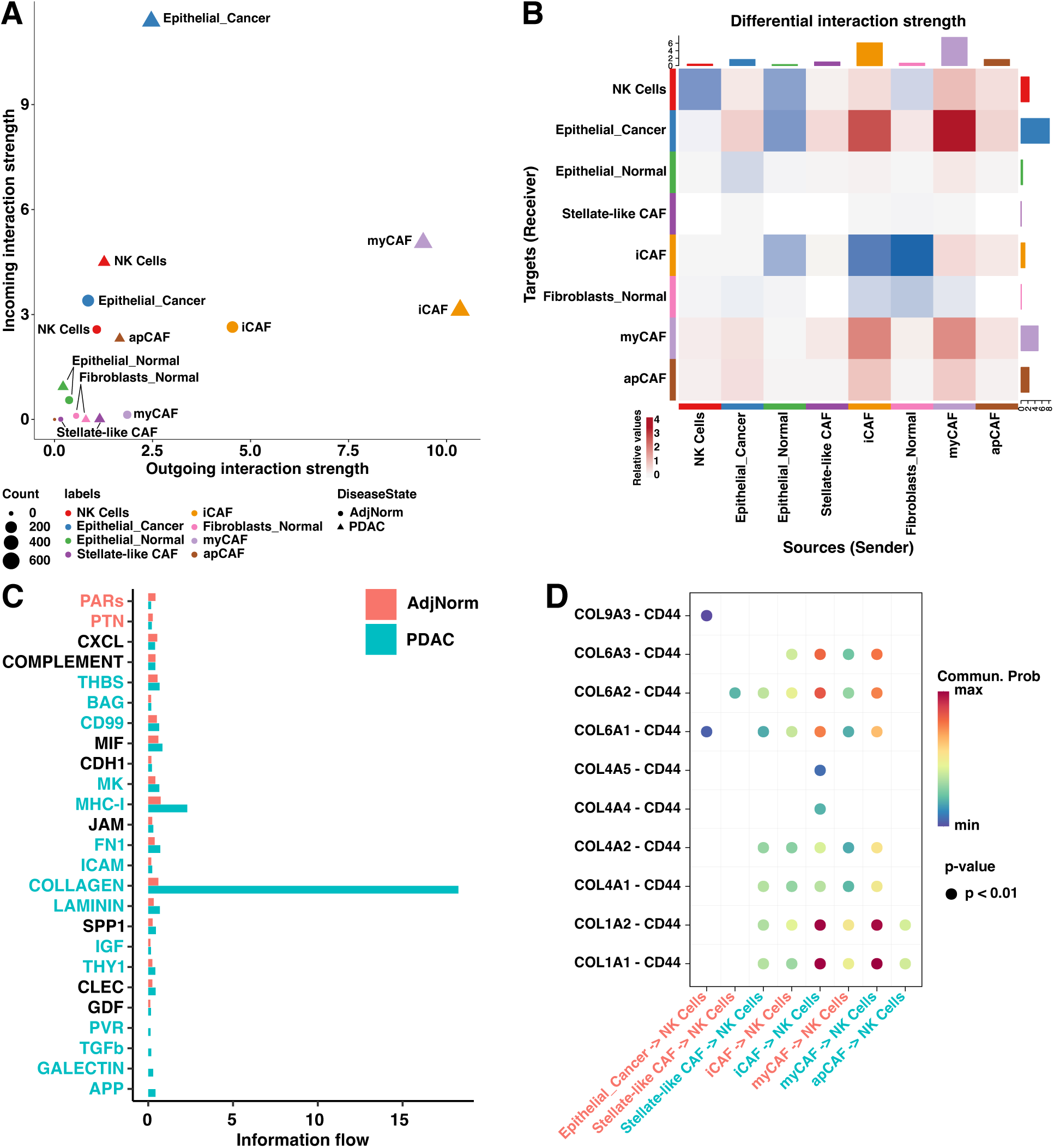
Cell-cell communication analysis implicates CD44a stromal checkpoint mediating NK- ECM interactions. **A.** Incoming and outgoing interaction strength for each cell population grouped by AdjNorm and PDAC disease states. **B.** Heatmap of differential interaction strengths of target (y-axis) and source (x-axis) cells with bar plots displaying the absolute values of interactions. **C.** Communication probabilities of signaling pathways targeting NK cells between AdjNorm and PDAC disease states as measured by information flow (pathway legend: pink = AdjNorm-specific, teal = PDAC-specific, black = equally specific). **D.** Significant ligand-receptor pairs from the COLLAGEN signaling pathways targeting NK cells upregulated in PDAC compared to AdjNorm (interaction legend (x-axis): pink = AdjNorm, teal = PDAC).

Among enriched signaling pathways, collagen signaling showed the largest PDAC-specific increase, driven by upregulation of multiple collagen isoforms (**Fig. 3C**). Within this network, NK cells were predicted to receive collagen-derived signals through CD44 (**Fig. 3D**), a principal receptor mediating cell–matrix adhesion on leukocytes^45^. Differential expression analysis confirmed upregulated CD44 expression in NK cells within the PDAC TME, concurrent with transcriptional upregulation of collagen genes in malignant epithelial cells, iCAFs, and myCAFs (**Fig. 3D, Supplementary Fig. S9**).

Together, these findings implicate the collagen–CD44 axis as a critical CAF-to-NK communication route that reinforces stromal confinement of cytotoxic lymphocytes. This represents a stromal checkpoint mechanism in which NK cells are sequestered within collagen-rich regions. We hypothesize that this CAF-NK cell signaling pathway limits the migration of NK cells toward tumor targets. Hence, disrupting CD44–collagen interactions could restore NK-cell motility and enhance antitumor function in PDAC.

### CD44 blockade enhances NK cell invasion *in vitro*

Given that NK cells in the PDAC TME primarily engage the collagen pathway through CD44 (**Figs. 3C-D**), we hypothesized that CD44-mediated interactions with ECM components physically restrict NK cell infiltration by anchoring them within the stroma. To test this, we first confirmed CD44 expression on human donor NK cells by flow cytometry (**Supplementary Fig. S10**). We then evaluated NK cell invasion through basement-membrane and collagen-containing matrices *in vitro.* In a 2D Transwell assay, membranes were coated with Matrigel (rich in ECM components^46^), and NK cells were incubated with or without a CD44-neutralizing antibody (α-CD44) (**Fig. 4A**). CD44 blockade significantly enhanced NK cell invasion in three of four human donor NK cell samples (**Figs. 4B-E**).

**Figure 4:**
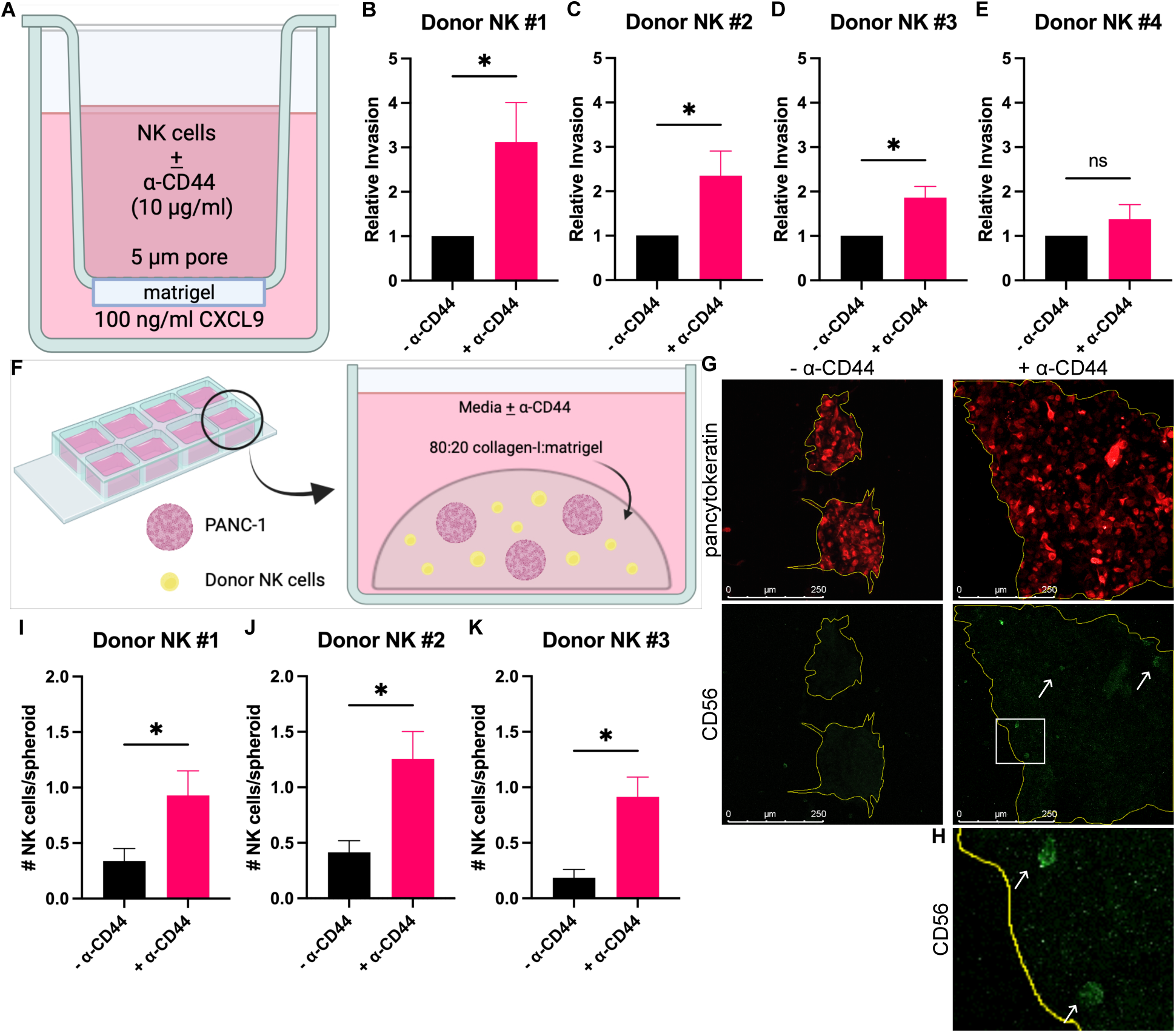
CD44 blockade enhances NK cell invasion *in vitro*. **A.** Schematic for Transwell invasion assays. Average relative invasion (± SEM) of **B.** Donor NK #1 (n = 9), **C.** Donor NK #2 (n = 12), **D.** Donor NK #3 (n = 6), and **E.** Donor NK #4 (n = 10) upon CD44 neutralization. *p < 0.05 as determined by Wilcoxon matched-pairs signed rank test. **F.** Schematic of spheroid invasion assay. **G.** Representative 20X IF images of outlined (yellow) PANC-1 spheroids (Pan-cytokeratin; red) embedded with NK cells (CD56; green), pointed out with white arrows, treated with or without α-CD44. Scale bar = 250 µm. **H.** Zoomed inset of NK cells in the α-CD44 treatment group in **G** (white box). Average number of NK cells per PANC-1 spheroid (± SEM) from **I.** Donor NK #1 (- α-CD44, n = 53; + α-CD44, n = 42), **J.** Donor NK #2 (- α-CD44, n = 29; + α-CD44, n = 43), and **K.** Donor NK #3 (- α-CD44, n = 27; + α-CD44, n = 47); n = # of spheroids analyzed; \**p* < 0.05 as determined by Wilcoxon matched-pairs signed rank test.

To validate these findings in a more physiological context, we employed a 3D *in vitro* model using a collagen-rich tumor spheroid assay (**Fig. 4F**). Human PANC-1 spheroids were embedded with donor NK cells in a Collagen I–Matrigel matrix and cultured for 24 hours with or without α-CD44. Immunofluorescence imaging showed increased NK-cell penetration into PANC-1 spheroids upon CD44 blockade across three donor NK cells (**Figs. 4G-K**, **Supplementary Fig. S11**). To determine whether this enhanced invasion involved CD44-dependent modulation of extracellular matrix degradation^47, 48^, we analyzed matrix metalloproteinase (MMP) gene expression in human donor NK cells treated with or without α-CD44 for 1 or 4 hours^47, 48^. No significant differences were observed (**Supplementary Fig. S12, Supplementary Table S5**), strongly suggesting that CD44 neutralization facilitates NK cell invasion primarily by disrupting adhesion to collagen, rather than altering proteolytic activity.

Together, these results demonstrate that CD44 functions as an inhibitory anchor for NK cells within the collagen-rich PDAC stroma. Blocking CD44–collagen binding restores NK-cell mobility and invasive capacity, providing a mechanistic rationale for targeting stromal checkpoints to enhance lymphocyte infiltration in desmoplastic tumors.

## Discussion

Recent reports characterizing immune-stromal interactions in PDAC using single-cell technologies have characterized the stromal heterogeneity and impact of CAFs on the PDAC TME^8–18^. However, the impact of stromal architecture on CAF, tumor cell, and leukocyte interactions and leukocyte infiltration in malignant ductal regions, the critically important tumor-immune interface, remains unexplored. Here, we define CAF-stratified ductal spatial architecture as a key organizational feature of the PDAC TME that dictates leukocyte access to malignant epithelium. Utilizing an approach that combines IMC, mIHC, and scRNA-seq to map periductal cellular interactions, we have demonstrated that periductal fibroblast organization is a critical determinant of leukocyte infiltration in malignant ductal regions. IMC and mIHC analyses collectively revealed that fibroblast abundance inversely correlated with immune accessibility, with NK cells being the leukocyte subset most sensitive to fibrous density. These data establish periductal fibroblast composition, not total fibrosis, as the dominant structural determinant of immune exclusion in PDAC.

Cell-to-cell communication analysis revealed that CAFs function as the dominant senders of signals within the PDAC TME, engaging both epithelial and NK cells through collagen-mediated pathways. The collagen-CD44 axis emerged as the principal CAF-to-NK signaling pathway, positioning CD44 as a stromal checkpoint that restricts leukocyte motility and reinforces fibroblast-mediated immune exclusion at malignant ducts. Functionally, CD44 blockade restored NK cell infiltration through collagen-rich matrices without altering MMP expression, indicating that neutralizing CD44–collagen adhesion, not ECM degradation, overcomes such barriers.

In PDAC, stromal elements, including CAF-derived chemokines (e.g., CXCL12)^49^ and ECM-derived ligands, such as fibronectin that engage inhibitory receptors like ILT3^50^, have been shown to modulate immune suppression within the TME. For example, elevated stromal CXCL12 correlates with the exclusion of cytotoxic T cells from tumor nests, while fibronectin engagement of the inhibitory receptor ILT3 on intra-tumoral myeloid cells induces a suppressive phenotype, collectively demonstrating that stromal components can directly regulate immune positioning and activation. Together, these findings indicate that the desmoplastic stroma functions not only as a physical barrier but also as an active regulatory element, acting as a “stromal checkpoint,” in which CAF-derived and ECM-mediated signals restrict leukocyte access and engagement with malignant epithelial cells. Our data extend this framework by identifying impaired immune trafficking as a key regulatory mechanism mediated through CD44–ECM interactions. Given that broad stromal ablation and ECM degradation have proven unsuccessful in restoring intra-tumoral leukocyte functionality or limiting tumor growth preclinically and clinically^51, 52^, our data highlight the potential of instead targeting stromal checkpoints, such as CD44, to enhance NK cell motility and infiltration, thus improving the efficacy of NK cell-dependent immunotherapies in PDAC.

In summary, our findings establish fibroblast composition as a central determinant of PDAC tissue architecture and immune accessibility. Focusing on malignant ductal regions, we define CAF-stratified ductal spatial architecture as the structural basis of immune exclusion and identify CD44 as a molecular mediator of this stromal checkpoint. High-CAF ducts promote epithelial cell proliferation while restricting lymphocyte access, particularly NK cells, whereas low-CAF ducts preserve effector function and cytolytic ability. Thus, targeting stromal checkpoints, in combination with immunotherapy and KRAS inhibitors, presents an opportunity to increase leukocyte infiltration irrespective of ECM degradation in PDAC and other solid tumors.

## Methods

### Tissue samples, antibody validation, metal-conjugation, and imaging mass cytometry (IMC)

The Lombardi Comprehensive Cancer Center (LCCC) Histopathology & Tissue Shared Resource (HTSR) provided formalin-fixed, paraffin-embedded (FFPE) human PDAC and spleen tissue samples. For IMC, we used a LCCC-curated FFPE human pancreas tissue microarray (TMA) slide series containing 2 mm cores from normal pancreas, pancreatitis, pancreatic neuroendocrine tumors, intraductal papillary mucinous neoplasm, pancreatic mucinous adenocarcinoma, PDAC, colon adenocarcinoma, testicle, tonsil, and placenta samples, and PDAC cell lines. Information for all TMA samples are available at doi: 10.5281/zenodo.10582038. The IMC study was restricted to 44 cores from 21 previously untreated PDAC patients who underwent Whipple resections or distal pancreatectomies (**Supplementary Table S2**).

For IMC, we designed a metal-conjugated antibody panel. Carrier-free antibodies not commercially metal-conjugated were validated by IF in PDAC or spleen tissue (**Supplementary Table S1**). Antibodies with expected expression patterns were metal-conjugated using MaxPar X8 Multimetal Labeling Kits (Standard Bio Tools (SBI); (**Supplementary Table S1**). Metal-conjugation was validated by cytometry by time-of-flight (CyTOF) (Hyperion Imaging System; SBI) at the LCCC Flow Cytometry Shared Resource (FCSR). Metal-conjugated antibody dilutions were optimized in the pancreas TMA prior to final IMC staining at the HTSR. Antigen retrieval and staining was performed (SBI) at the LCCC HTSR, while image acquisition was performed at the LCCC Mass Spectrometry and Analytical Pharmacology Shared Resource. A maximum of 2.25 mm^2^ regions of interest (ROIs) on TMA cores were selected, laser-ablated and analyzed by CyTOF to quantitate the metal-conjugated antibodies per ROI, generating multiplex images^53^. All data were collected using the Hyperion Imaging System (SBI) and qualitatively validated in MCD Viewer (v1.0.560.6; SBI) and QuPath softwares (RRID:SCR_018257).

### IMC image processing, data transformation, and single-cell clustering

Raw IMC data was processed following the widely used ImcSegmentationPipeline^54^ (https://github.com/BodenmillerGroup/ImcSegmentationPipeline). In the Ilastik software (RRID:SCR_015246), DNA, E-cadherin, pan-cytokeratin, vimentin, and CD45RO markers were used to pixel-train and generate cell segmentation probability maps for nuclear, cytoplasmic, cell membrane, and background areas in all IMC images, which were imported into CellProfiler to generate cell segmentation masks to then extract single-cell protein expression data using histoCAT^28^. 412,862 cells were identified across 44 images from 21 PDAC patient samples. Extracted IMC-derived single cell data were imported into R (v4.2.1) for downstream computational analyses. histoCAT-derived single cell data files generated are available at doi: 10.5281/zenodo.10582038. All scripts adapted from this pipeline for downstream analyses are available at https://github.com/Weiner-Lab/NaturalKillerCells_PDAC. Single cell protein expression data was log-transformed, re-integrated, and clustered using Seurat-based analysis^27^. As single cell expression may contain “lateral spillover” into neighboring single cells as a result of cell segmentation^54^, we therefore used a supervised clustering approach to phenotype cells based on canonical cellular morphology markers (αSMA, CD3, CD4, CD8α, CD11b, CD16, CD20, CD45RO, CD56, CD68, E-cadherin, FAP, FOXP3, NKG2D, NKp44, pan-cytokeratin, vimentin), we found that a clustering resolution = 0.35 provided distinct immune, fibroblast, and epithelial cell populations. Each cell population was subsetted and re-clustered to identify the functional state of subclusters using the following markers: CD69, CD107α, cleaved-caspase 3, collagen-I, granzyme B, HLA-A/B/C, HLA-A1, IFNγ, Ki67, MICA, PCNA, perforin, PD-1, PD-L1, and TGFβ. All cellular subclusters were reintegrated and annotated based on functional classification: active subpopulations were solely immune cell based and were characterized by high expression of known immune cell activation markers, CD69, CD107α, and/or IFNγ; proliferative cell populations were characterized by high expression of proliferation markers Ki67 and/or PCNA; and inhibitory subpopulations were characterized by high TGFβ expression, which is known to be immune suppressive. Subclustering identified 29 unique cell populations, which were validated visually using the Cytomapper R package and the MCD Viewer software. Seurat objects containing all the PDAC single cell metadata is available at doi: 10.5281/zenodo.10582038 and were converted to SingleCellExperiment^54^ objects for downstream spatial analyses.

### IMC spatial analysis

We generated knn-based interaction graphs for the 15-nearest neighbors and used k-means clustering to group cells into 8 distinct cellular neighborhoods using the SpatialExperiment and imcRtools R packages^54^. Using the imcRtools package, permutation test-based cell-cell interaction analysis identified significant interactions or avoidances between cell populations. Using FDR analysis, cell-cell interactions with *p* < 0.01 were considered statistically significant. Based on Euclidean distance metadata, distance and network analyses were performed, as previously described^33^. Seurat’s CellSelector function^27^ was used to select and quantify cells in epithelial-ductal ROIs of spatial images. From CellSelector, percentages of cell populations in each epithelial-ductal ROI were quantified, and epithelial-ductal ROIs were stratified in quartiles based on fibroblast abundance (Q1 = low fibroblast quartile, Q4 = high fibroblast quartile). Mean total expression of all markers was quantified in representative epithelial-ductal ROIs of pseudoimages using MCD Viewer. Cell-based expression in epithelial-ductal ROIs was performed using Seurat.

### Multiplex immunohistochemistry (mIHC)

Sections of 5μm thickness were cut from FFPE tissue blocks containing PDAC tumor samples. Patient samples were from 10 males and 7 females with age ranges between 45-88, and had tumor diameters ranging from 0.1-8 cm (**Supplementary Table S3**). Samples were stained for CD56, CD16, CD335, panCK, αSMA, and CD68 (**Supplementary Table S4**).

The slides were baked at 60°, deparaffinized in xylene, rehydrated, washed in distilled water and incubated with 10% neutral buffered formalin (NBF) for an additional 20 minutes to increase tissue-slide retention. Epitope retrieval/microwave treatment (MWT) for all antibodies was performed by boiling slides in the respective Epitope Retrieval buffer (ER1, pH6 or ER2, pH9; Leica Biosystems AR9961 or AR9640, respectively). Protein blocking was performed using antibody diluent/blocking buffer (Akoya, ARD1001EA) for 10 minutes at room temperature. Primary antibody/OPAL dye pairings, staining order and incubation conditions are listed in the table below. The staining was performed using Leica Bond autostainer. Sections were counterstained with spectral DAPI (Akoya FP1490) for 5 min and mounted with ProLong Diamond Antifade (ThermoFisher, P36961) using StatLab #1 coverslips (CV102450). The order of antibody staining and the antibody/OPAL pairing was predetermined using general guidelines and the particular biology of the panel. General guidelines include spectrally separating co-localizing markers and separating spectrally adjacent dyes. Multiplex IHC was optimized by first performing single-plex IHC with the chosen antibody/OPAL dye pair to optimize signal intensity values and proper cellular expression, followed by optimizing the full multiplex assay. Slides were scanned at 10X magnification using the Phenoimager Fusion system Automated Quantitative Pathology Imaging System (Akoya Biosciences). Whole slide scans were viewed with Phenochart (Phenochart 2.2.0, Akoya) which also allows for the selection of high-powered images at 20X (resolution of 0.5mm per pixel) for multispectral image capture. Multispectral images of each tissue specimen were captured in their entirety.

Multiplex staining and imaging were performed as previously described^55^, except tissue images were captured using Akoya Biosciences Inc.’s PhenoImager Fusion imaging platform at the LCCC HTSR. Images were exported as *.qptiff* files and visualized and processed using QuPath software (RRID:SCR_018257).

### mIHC image processing and analysis

mIHC images were processed using QuPath software (v0.6.0)^56^. Spectrally unmixed images were visualized and a ROI grid, with each ROI measuring 500 µm via 500 µm, was overlaid on each image. Cells within each ROI were then automatically segmented using the StarDist machine-learning based pipeline^57^, with the DAPI channel used to delineate nuclear staining, normalized percentile set at (1, 99), threshold set at 0.5, and pixel size set at 0.5 µm/pixel. After cell segmentation, duplicate channel training images were created for each marker to create an object classifier for each cell type. Under the direction of an experienced pathologist, an object classifier was trained for each marker. This was repeated for three separate images. Afterwards, object classifiers for each marker were then applied to each image for cell phenotyping. Percentage of cells was calculated by dividing cells detected for marker of interest by total cells in each ROI. All ROIs were then averaged to get a total value for each cell type of interest for each image. Only ROIs with at least 500 cells were analyzed, with biologically incompatible markers excluded from analysis.

### Single-cell RNA-sequencing analysis

To infer ligand-receptor (L-R) interactions in PDAC, we utilized a previously published human PDAC scRNAseq dataset^23^ and subsetted NK cells, fibroblasts, and epithelial cells. Malignant and normal epithelial cells were differentiated using CytoTRACE, a computational framework that predicts cellular differentiation states by measuring the number of genes expressed by each cell^41^. The CytoTRACE scores distribution was observed to be bi-modal and using a mixed-model, an ideal cut-off value separating the two distributions was determined to be 0.72 where scores greater than 0.72 indicated a malignant epithelial cells^58^ (**Supplementary Fig. S8**). Fibroblasts were subclustered into known subpopulations and we identified: antigen-presenting CAFs (apCAFs), inflammatory CAF (iCAFs), myofibroblasts (myCAFs), normal fibroblasts, and stellate-like fibroblasts^43^ (**Supplementary Fig. S8**). We then used CellChat, a manually curated protein database of approximately 2,000 L-R pairs^40^ to infer potential L-R interactions between epithelial, fibroblast, and NK cell populations. A Wilcox test with a Bonferroni correction from the FindMarkers Seurat function (v4.3) was used for differential gene expression analysis of CellChat-identified L-R genes. Genes with an adjusted-*P* < 0.01 were considered significantly differentially expressed. R scripts for applied CytoTRACE and CellChat analyses are available at https://github.com/Weiner-Lab/NaturalKillerCells_PDAC.

### Cell culture

Human PANC-1 PDAC cells and donor NK cell lines were cultured, as previously described^59, 60^, except donor NK cells received 0.5% v/v of IL-2. Healthy human donor NK cell lines, Donor NK #1, #2, and #4 were purchased (STEMCELL Technologies, Vancouver, Canada), and Donor NK #3 cells were isolated from a leukoreduction system (LRS) chamber (Inova Health System) and isolated via MojoSortTM Human NK Cell Isolation Kit (BioLegend, CAT#: 480053). Donor NK cells were stimulated with the engineered K562 cell line (RRID:CVCL_0004), K562-mb21-41BBL (courtesy of Dr. Dean Lee at Nationwide Children’s)^61^. K562-mb21-41BBL cells were cultured in supplemented RPMI (10% FBS, 1% penicillin/streptomycin). For NK cell stimulation, K562-mb21-41BBL cells were collected (5 minutes, 1,000 rpm, 4°C), resuspended in supplemented RPMI with mitomycin C (Sigma-Aldrich) (C_f_ = 0.01 mg/ml) and incubated for 4 hours (37°C). Pre-treated K562-mb21-41BBL cells were collected (5 minutes, 1,000 rpm, 4°C), resuspended in NK cell medium, then co-cultured with donor NK cells (E:T = 1:1); this process was repeated at least once per week. Donor NK cells aged day 1-50 and stimulated 1-3 days prior to *in vitro* assays were used. All cell lines were tested and determined to be free of mycoplasma.

### Flow cytometry

Cells were centrifuged (5 minutes, 1,000rpm, 4°C) washed twice (1X PBS), then resuspended in FACS staining buffer (1% BSA in 1X PBS). Human Fc block (BD Pharmigen, CAT#: 564219) then α-CD44-BV605 (BioLegend, CAT#: 103047) antibodies were added per manufacturer’s recommendations (30 minutes, 4°C). Cell pellets were washed twice (FACS staining buffer), resuspended in fixative (1% PFA in 1X PBS), then processed by the LCCC FCSR in a BD LSRFortessa Cell Analyzer (BD Biosciences). Analysis was performed using FlowJo (v10.8.1; RRID:SCR_008520).

### NK cell 2D invasion assay

The bottom of 5 µm Transwells (Corning) were coated with Matrigel (Corning) diluted 1:3 in NK cell medium. CXCL9 (C_f_ =100 ng/ml) (R&D Systems)-supplemented NK cell medium was added to the bottom of each well. Donor NK cells were added to Transwells with or without an α-CD44 neutralizing antibody (clone IM7; Bio X Cell; C_f_ = 10 µg/ml) and incubated for 4 hours (37°C). Transwells were removed, cells were stained with Trypan Blue Stain (Gibco) and manually counted on a hemocytometer. Control cells were plated without Transwells to count total cells to calculate percent invasion. Relative invasion is the percent of cells that invaded through the transwell normalized to untreated NK cell conditions. A Wilcoxon matched-pairs signed rank test was performed to determine statistical significance (GraphPad Prism v10; RRID:SCR_002798).

### NK cell 3D spheroid invasion assay

As previously described,^27^ PANC-1 cells were seeded in each well of a low-adhesion U-bottom plate, spun (5 minutes, 1,000 rpm, 4°C) and cultured overnight. Spheroids were embedded with donor NK cells in an 80:20 Collagen-I (Invitrogen): Matrigel (Corning) mixture in chamber wells (Lab-Tek). Embedded spheroids were treated with α-CD44 (clone IM7; Bio X Cell; C_f_ = 10 µg/ml) for 24 hours (37°C) prior to immunocytochemistry (IC). For IC, samples were fixed with 4% paraformaldehyde (PFA; 10 minutes, 25°C), washed with 1X PBS (5 minutes), then permeabilized with 0.5% Triton-X100 (10 minutes). Cells were washed once, blocked (1% BSA in 1X PBS, 30 minutes), then incubated with human Fc block (BD Pharmigen, CAT#: 564219). Samples were incubated with pan-cytokeratin-eFluor™ 660 (Invitrogen, CAT#: 50-9003-82) and CD56 (Invitrogen, CAT#: MA1-19129) antibodies overnight (25°C). An α-Mouse IgG2a-AF488 secondary antibody (Invitrogen, CAT#: A-21131; 1-4 hours, 25°C), cells were washed 3 times, incubated with DAPI (50 ng/ml; 10 minutes, 25°C), then washed twice. Coverslips were mounted with anti-fade mountant, then and dried. Z-stacks of spheroids were imaged using a Leica SP8 AOBS microscope at the LCCC Microscopy and Imaging Shared Resource. Z-stacks were ≤ 1 µm. Z-stack images were processed and using FIJI software^62^ (v2.14.0; RRID:SCR_002285) for manual NK cell quantitation into PANC-1 spheroids. Statistical significance was determined using a Wilcoxon matched-pairs signed rank test (GraphPad Prism v10).

### Effects of CD44 neutralization on MMP gene expression in human NK cells

Donor NK cell lines were incubated with or without α-CD44 (clone IM7; Bio X Cell; C_f_ = 10 µg/ml) for 1 and 4 hours. Cells were collected (5 minutes, 1,000 rpm, 4°C) and RNA was isolated using the PureLink™ RNA Mini Kit (Ambion). RNA concentration was measured using NanoDrop 8000 (ThermoFisher Scientific), then stored at −80°C. Diluted RNA (20 ng/µl) was treated with 10% v/v 10X DNase I Buffer and 10% v/v DNase I (Promega; 15 minutes, 25°C). 10% v/v of RQ1 RNase/DNase Stop Solution (Promega) and 10% v/v Oligo(dT) nucleotides (IDT) were added, then heat inactivated at 65°C (10 minutes) then at 4°C (5 minutes). cDNA was synthesized with the GoScript™ Reverse Transcriptase Kit (Promega). qPCR was performed using GoTaq qPCR Master Mix (Promega) with the StepOnePlus™ Real-Time PCR System (ThermoFisher Scientific). Gene expression was quantified (2^-ΔCт^; Ambion) using *HPRT* (Forward 3’→5’: CTTTCCTTGGTCAGGCAGTA, reverse 3’→5’: TGGCTTATATCCAACACTTCG) as an internal, endogenous control. Fold-change was calculated using the 2^-ΔΔCт^ method and a One-Way ANOVA, Kruskal-Wallis test was performed to determine statistical significance (GraphPad Prism v10).

## Ethics statement

The human pancreas TMA and PDAC slides used in this study were approved by the Georgetown University LCCC Biospecimen Use Committee.

## Supplementary Materials

Supplementary Figure S1: Spatial proteomic analysis of the human PDAC TME resolves PDAC cell populations.

Supplementary Figure S2: Subclustered cell populations of spatial proteomic resolved PDAC cell populations.

Supplementary Figure S3: Proportions of PDAC subclustered cell populations per cellular neighborhood.

Supplementary Figure S4: Pro-inflammatory NK cell and Macrophage Content is Inversely Correlated with αSMA+ Fibroblasts.

Supplementary Figure S5: PDAC epithelial-ductal ROI cell type abundance.

Supplementary Figure S6: PDAC epithelial-ductal ROI cell subcluster abundance.

Supplementary Figure S7: NK cells retain expression of activator markers in fibroblast-rich periductal regions.

Supplementary Figure S8: Subclustering of epithelial and fibroblast populations in human PDAC scRNAseq samples.

Supplementary Figure S9: Fibroblast and epithelial cell populations exhibit differential expression of CellChat-based collagen signaling genes.

Supplementary Figure S10: Human donor NK cells express CD44.

Supplementary Figure S11: CD44 neutralization increases human donor NK cell invasion into PANC-1 PDAC spheroids.

Supplementary Figure S12: CD44 neutralization does not affect expression of MMP genes in human donor NK cells.

Supplementary Table S1: IMC antibody panel and metal assignments.

Supplementary Table S2: PDAC patient information for IMC.

Supplementary Table S3: PDAC patient information for multiplex IF (Vectra®).

Supplementary Table S4: Antibodies and fluorescent dyes for multiplex IF (Vectra®).

Supplementary Table S5: MMP genes primer sequences.

## Acknowledgments

We would like to thank the following from the Georgetown University Lombardi Comprehensive Cancer Center’s Shared Resources (SR) (partially supported by NIH/NCI P30CA51008 grant): Dr. Karen Creswell, Dan Xu, and Zhinuo Jiang at the Flow Cytometry and Cell Sorting SR (partially supported NIH S10OD016213 grant), Aaron Rozeboom, Bin Li, and Dr. Susree Modepalli at the Histopathology and Tissue SR, Dr. Junfeng Ma at the Mass Spectrometry and Analytical Pharmacology and the Biostatistics SR for support with performing metal-conjugation, IMC, and multiplex immunohistochemistry, and with providing guidance on spatial analyses. We would also like to thank the Tissue Culture and Biobanking and Microscopy and Imaging Shared Resources for support on *in vitro* assays. We also thank Ahmed Elhossiny and the Pasca Di Magliano lab at the University of Michigan for providing us with the scRNAseq dataset for analyses described here. We also thank Drs. Andrew Quong and Eric Swanson at Standard BioTools Inc. for assistance with the IMC antibody panel design and its cell segmentation pipeline, and Dr. Ludmila Danilova at Johns Hopkins University for advice on computational analyses. Schematics were created with BioRender.com.

## Author contributions

Z.X.M., A.A.L., and L.M.W. conceived the idea, designed the study, obtained funding, and wrote the manuscript. Z.X.M. and A.A.L. conducted experiments and analyzed the data. Z.X.M., A.A.L., R.K.S., A.D., S.A., and W.J.H. performed or assisted with computational analyses. Z.X.M., R.K.S., and S.A. wrote R scripts and generated the computational figures. B.H. and A.D. assisted with data collection and tissue pathology of human tissue samples. L.W. and M.C. assisted in performing 2D *in vitro* invasion experiments. M.C. assisted with RT-qPCR, flow cytometry assays, and 3D *in vitro* invasion experiments. I.P. and S.H. assisted in IMC data collection. W.J.H., E.J.F., M.A., and S.A.J. assisted with experimental design. M.A., E.J.F, and L.M.W. edited the manuscript. All authors critically reviewed and approved the manuscript.

## Competing interests

Z.X.M. is currently employed by AstraZeneca. E.J.F. was on the scientific advisory board of Resistance Bio and a consultant for Mestag Therapeutics. L.M.W. sits on the Scientific Advisory Boards of Celldex Therapeutics, Bobcat Bio, Kuiper and Cytomx Therapeutics, is an advisor for Fortress Biotech and is founder and Chair of the Board of PushCART Therapeutics.

## Code availability statement

Code used to perform IMC and scRNAseq analyses and generate the figures is available at https://github.com/Weiner-Lab/NaturalKillerCells_PDAC.

## Data transparency statement

Code used to perform IMC and scRNAseq analyses and generate the figures is available at https://github.com/Weiner-Lab/NaturalKillerCells_PDAC. Raw .mcd files for all cores of the pancreas tissue microarray imaged by IMC and IMC-derived datasets, including histoCAT-derived single cell files, and Seurat, SingleCellExperiment, Cytomapper and distance data matrix R objects, to reproduce figures are deposited at doi: 10.5281/zenodo.10582038. Source Data are provided with this paper. The scRNAseq data was provided to us by the Pasca Di Magliano lab at the University of Michigan, which was previously published at doi: 10.1038/s43018-020-00121-4. The authors declare that all other data supporting the findings of this study are available within the paper or its supplementary information files. All other relevant data are available from the corresponding author upon reasonable request.

## Funding

This work is supported by grants from the National Institutes of Health, National Cancer Institute T32CA009686 (Training Grant in Tumor Biology, Anna Tate Riegel), F31CA261125 (Z.X.M.), F30CA294875 (A.A.L.), U24CA284156 (E.J.F.), and P30CA51008 (L.M.W), Achievement Reward for College Scientists (ARCS) Foundation Scholar (Z.X.M.), and the Lustgarten Foundation (E.J.F and W.J.H.).

## Notes

https://www.zenodo.org/10.5281/zenodo.10582038

### Summary of Updates

The current submission represents a conceptually re-envisioned study that diverges substantially from the earlier version in focus, scope, and mechanistic insight. In the prior submission, our work centered on the spatial distribution and epithelial interactions of natural killer (NK) cells within the pancreatic ductal adenocarcinoma (PDAC) microenvironment. Since then, we have fundamentally reframed the study to address a broader and more impactful question: how the malignant ductal-fibrotic interface governs immune exclusion through a CD44-dependent stromal checkpoint. Major conceptual and experimental advances include: Reframed focus from NK cells to pan-leukocyte spatial organization at the malignant ductal regions -We shifted the emphasis from a single immune subset (NK cells) to a comprehensive analysis of all leukocyte populations within PDAC epithelial-ductal regions. -This reframing positions the study within a new conceptual framework, the periductal fibroblast density as a defining determinant of immune accessibility and epithelial proliferation. Identification of a stromal checkpoint governing immune access. Through integrated spatial and single-cell analyses, we defined a "stromal checkpoint" that restricts immune cell access to malignant ducts. Within this framework, collagen-CD44 adhesion emerged as a key mechanistic axis mediating immune cell retention in fibroblast-rich zones. Functional 3D spheroid assays demonstrated that disrupting this interaction enhances immune motility independently of ECM degradation, establishing a new conceptual model in which stromal architecture--not immune cell exhaustion--acts as the rate-limiting barrier to infiltration. Validation of IMC analysis We validated our IMC findings, by performing Vectra multiplex immunofluorescence (mfIHC) on an independent cohort of 17 PDAC tissues, confirming the inverse correlation between periductal fibroblast abundance and pro-inflammatory immune infiltration. Conceptual and structural overhaul of the manuscript -The new version has been retitled, reorganized, and rewritten to reflect this paradigm shift, from a descriptive NK-cell study to a mechanistically grounded analysis of stromal architecture as an immune barrier. - The manuscript now defines a broadly applicable mechanism of immune exclusion relevant to tumor immunology, microenvironmental biology, and translational oncology.

## References

(1) Siegel, R. L.; Giaquinto, A. N.; Jemal, A. Cancer statistics, 2024. CA: a cancer journal for clinicians 2024, 74 (1), 12–49. DOI: 10.3322/caac.21820.

(2) Royal, R. E.; Levy, C.; Turner, K.; Mathur, A.; Hughes, M.; Kammula, U. S.; Sherry, R. M.; Topalian, S. L.; Yang, J. C.; Lowy, I. Phase 2 trial of single agent Ipilimumab (anti-CTLA-4) for locally advanced or metastatic pancreatic adenocarcinoma. Journal of immunotherapy 2010, 33 (8), 828–833.

(3) Brahmer, J. R.; Tykodi, S. S.; Chow, L. Q.; Hwu, W.-J.; Topalian, S. L.; Hwu, P.; Drake, C. G.; Camacho, L. H.; Kauh, J.; Odunsi, K. Safety and activity of anti–PD-L1 antibody in patients with advanced cancer. New England journal of medicine 2012, 366 (26), 2455–2465.

(4) O’Reilly, E. M.; Oh, D. Y.; Dhani, N.; Renouf, D. J.; Lee, M. A.; Sun, W.; Fisher, G.; Hezel, A.; Chang, S. C.; Vlahovic, G.;, et al. Durvalumab With or Without Tremelimumab for Patients With Metastatic Pancreatic Ductal Adenocarcinoma: A Phase 2 Randomized Clinical Trial. JAMA Oncol 2019, 5 (10), 1431–1438. DOI: 10.1001/jamaoncol.2019.1588 From NLM PubMed-not-MEDLINE.

(5) Karamitopoulou, E. Tumour microenvironment of pancreatic cancer: immune landscape is dictated by molecular and histopathological features. British journal of cancer 2019, 121 (1), 5–14. DOI: 10.1038/s41416-019-0479-5.

(6) Muller, M.; Haghnejad, V.; Schaefer, M.; Gauchotte, G.; Caron, B.; Peyrin-Biroulet, L.; Bronowicki, J.-P.; Neuzillet, C.; Lopez, A. The immune landscape of human pancreatic ductal carcinoma: key players, clinical implications, and challenges. Cancers 2022, 14 (4), 995. DOI: 10.3390/cancers14040995.

(7) Ferdek, P. E.; Jakubowska, M. A. Biology of pancreatic stellate cells—more than just pancreatic cancer. Pflügers Archiv-European Journal of Physiology 2017, 469, 1039–1050.

(8) Khaliq, A. M.; Rajamohan, M.; Saeed, O.; Mansouri, K.; Adil, A.; Zhang, C.; Turk, A.; Carstens, J. L.; House, M.; Hayat, S. Spatial transcriptomic analysis of primary and metastatic pancreatic cancers highlights tumor microenvironmental heterogeneity. Nature genetics 2024, 56 (11), 2455–2465.

(9) Elyada, E.; Bolisetty, M.; Laise, P.; Flynn, W. F.; Courtois, E. T.; Burkhart, R. A.; Teinor, J. A.; Belleau, P.; Biffi, G.; Lucito, M. S. Cross-species single-cell analysis of pancreatic ductal adenocarcinoma reveals antigen-presenting cancer-associated fibroblasts. Cancer discovery 2019, 9 (8), 1102–1123.

(10) Xu, Y.; Wang, X.; Li, Y.; Mao, Y.; Su, Y.; Mao, Y.; Yang, Y.; Gao, W.; Fu, C.; Chen, W. Multimodal single cell-resolved spatial proteomics reveal pancreatic tumor heterogeneity. Nature Communications 2024, 15 (1), 10100.

(11) Sussman, J. H.; Kim, N.; Kemp, S. B.; Traum, D.; Katsuda, T.; Kahn, B. M.; Xu, J.; Kim, I.-K.; Eskandarian, C.; Delman, D. Multiplexed imaging mass cytometry analysis characterizes the vascular niche in pancreatic cancer. Cancer research 2024, 84 (14), 2364–2376.

(12) Yousuf, S.; Qiu, M.; von Voithenberg, L. V.; Hulkkonen, J.; Macinkovic, I.; Schulz, A. R.; Hartmann, D.; Mueller, F.; Mijatovic, M.; Ibberson, D. Spatially resolved multi-omics single-cell analyses inform mechanisms of immune dysfunction in pancreatic cancer. Gastroenterology 2023, 165 (4), 891–908. e814.

(13) Cui Zhou, D.; Jayasinghe, R. G.; Chen, S.; Herndon, J. M.; Iglesia, M. D.; Navale, P.; Wendl, M. C.; Caravan, W.; Sato, K.; Storrs, E. Spatially restricted drivers and transitional cell populations cooperate with the microenvironment in untreated and chemo-resistant pancreatic cancer. Nature genetics 2022, 54 (9), 1390–1405.

(14) Kim, S.; Leem, G.; Choi, J.; Koh, Y.; Lee, S.; Nam, S.-H.; Kim, J. S.; Park, C. H.; Hwang, H. K.; Min, K. I. Integrative analysis of spatial and single-cell transcriptome data from human pancreatic cancer reveals an intermediate cancer cell population associated with poor prognosis. Genome medicine 2024, 16 (1), 20.

(15) Liudahl, S. M.; Betts, C. B.; Sivagnanam, S.; Morales-Oyarvide, V.; Da Silva, A.; Yuan, C.; Hwang, S.; Grossblatt-Wait, A.; Leis, K. R.; Larson, W. Leukocyte heterogeneity in pancreatic ductal adenocarcinoma: phenotypic and spatial features associated with clinical outcome. Cancer discovery 2021, 11 (8), 2014–2031.

(16) Ene-Obong, A.; Clear, A. J.; Watt, J.; Wang, J.; Fatah, R.; Riches, J. C.; Marshall, J. F.; Chin-Aleong, J.; Chelala, C.; Gribben, J. G.;, et al. Activated pancreatic stellate cells sequester CD8+ T cells to reduce their infiltration of the juxtatumoral compartment of pancreatic ductal adenocarcinoma. Gastroenterology 2013, 145 (5), 1121–1132. DOI: 10.1053/j.gastro.2013.07.025 From NLM Medline.

(17) Bell, A. T.; Chianchiano, P.; Hirose, K.; Salas-Escabillas, D.; Zucha, D. M.; Mitchell, J. T.; Danilova, L.; Damanakis, A. I.; Tandurella, J. A.; Johnson, J. Spatial profiling of human pancreatic ductal adenocarcinoma reveals molecular alterations associated with venous invasion. Science Translational Medicine 2025, 17 (817), eady7524.

(18) Bell, A. T.; Mitchell, J. T.; Kiemen, A. L.; Lyman, M.; Fujikura, K.; Lee, J. W.; Coyne, E.; Shin, S. M.; Nagaraj, S.; Deshpande, A. PanIN and CAF transitions in pancreatic carcinogenesis revealed with spatial data integration. Cell Systems 2024, 15 (8), 753–769. e755.

(19) Biffi, G.; Oni, T. E.; Spielman, B.; Hao, Y.; Elyada, E.; Park, Y.; Preall, J.; Tuveson, D. A. IL1-induced JAK/STAT signaling is antagonized by TGFβ to shape CAF heterogeneity in pancreatic ductal adenocarcinoma. Cancer discovery 2019, 9 (2), 282–301.

(20) Oh, K.; Yoo, Y. J.; Torre-Healy, L. A.; Rao, M.; Fassler, D.; Wang, P.; Caponegro, M.; Gao, M.; Kim, J.; Sasson, A. Coordinated single-cell tumor microenvironment dynamics reinforce pancreatic cancer subtype. Nature communications 2023, 14 (1), 5226.

(21) Niu, N.; Shen, X.; Wang, Z.; Chen, Y.; Weng, Y.; Yu, F.; Tang, Y.; Lu, P.; Liu, M.; Wang, L. Tumor cell-intrinsic epigenetic dysregulation shapes cancer-associated fibroblasts heterogeneity to metabolically support pancreatic cancer. Cancer Cell 2024, 42 (5), 869–884. e869.

(22) Ligorio, M.; Sil, S.; Malagon-Lopez, J.; Nieman, L. T.; Misale, S.; Di Pilato, M.; Ebright, R. Y.; Karabacak, M. N.; Kulkarni, A. S.; Liu, A. Stromal microenvironment shapes the intratumoral architecture of pancreatic cancer. Cell 2019, 178 (1), 160–175. e127.

(23) Steele, N. G.; Carpenter, E. S.; Kemp, S. B.; Sirihorachai, V. R.; The, S.; Delrosario, L.; Lazarus, J.; Amir, E.-a. D.; Gunchick, V.; Espinoza, C. Multimodal mapping of the tumor and peripheral blood immune landscape in human pancreatic cancer. Nature cancer 2020, 1 (11), 1097–1112. DOI: 10.1038/s43018-020-00121-4.

(24) Chang, Q.; Ornatsky, O. I.; Siddiqui, I.; Loboda, A.; Baranov, V. I.; Hedley, D. W. Imaging mass cytometry. Cytometry part A 2017, 91 (2), 160–169. DOI: 10.1002/cyto.a.23053.

(25) Butler, A.; Hoffman, P.; Smibert, P.; Papalexi, E.; Satija, R. Integrating single-cell transcriptomic data across different conditions, technologies, and species. Nature biotechnology 2018, 36 (5), 411–420.

(26) Stuart, T.; Butler, A.; Hoffman, P.; Hafemeister, C.; Papalexi, E.; Mauck, W. M.; Hao, Y.; Stoeckius, M.; Smibert, P.; Satija, R. Comprehensive integration of single-cell data. cell 2019, 177 (7), 1888–1902. e1821.

(27) Hao, Y.; Hao, S.; Andersen-Nissen, E.; Mauck, W. M.; Zheng, S.; Butler, A.; Lee, M. J.; Wilk, A. J.; Darby, C.; Zager, M. Integrated analysis of multimodal single-cell data. Cell 2021, 184 (13), 3573–3587. e3529. DOI: 10.1016/j.cell.2021.04.048.

(28) Schapiro, D.; Jackson, H. W.; Raghuraman, S.; Fischer, J. R.; Zanotelli, V. R.; Schulz, D.; Giesen, C.; Catena, R.; Varga, Z.; Bodenmiller, B. histoCAT: analysis of cell phenotypes and interactions in multiplex image cytometry data. Nature methods 2017, 14 (9), 873–876. DOI: 10.1038/nmeth.4391.

(29) Satija, R.; Farrell, J. A.; Gennert, D.; Schier, A. F.; Regev, A. Spatial reconstruction of single-cell gene expression data. Nature biotechnology 2015, 33 (5), 495–502.

(30) Hafemeister, C.; Satija, R. Normalization and variance stabilization of single-cell RNA-seq data using regularized negative binomial regression. Genome biology 2019, 20 (1), 296. DOI: 10.1186/s13059-019-1874-1.

(31) Schürch, C. M.; Bhate, S. S.; Barlow, G. L.; Phillips, D. J.; Noti, L.; Zlobec, I.; Chu, P.; Black, S.; Demeter, J.; McIlwain, D. R. Coordinated cellular neighborhoods orchestrate antitumoral immunity at the colorectal cancer invasive front. Cell 2020, 182 (5), 1341–1359. e1319. DOI: 10.1016/j.cell.2020.07.005.

(32) Gocher, A. M.; Workman, C. J.; Vignali, D. A. Interferon-γ: teammate or opponent in the tumour microenvironment? Nature Reviews Immunology 2022, 22 (3), 158–172.

(33) Ho, W. J.; Zhu, Q.; Durham, J.; Popovic, A.; Xavier, S.; Leatherman, J.; Mohan, A.; Mo, G.; Zhang, S.; Gross, N. Neoadjuvant cabozantinib and nivolumab convert locally advanced hepatocellular carcinoma into resectable disease with enhanced antitumor immunity. Nature Cancer 2021, 2 (9), 891–903. DOI: 10.1038/s43018-021-00234-4.

(34) Lee, H.; Ferguson, A. L.; Quek, C.; Vergara, I. A.; Pires daSilva, I.; Allen, R.; Gide, T. N.; Conway, J. W.; Koufariotis, L. T.; Hayward, N. K. Intratumoral CD16+ macrophages are associated with clinical outcomes of patients with metastatic melanoma treated with combination anti-PD-1 and anti-CTLA-4 therapy. Clinical Cancer Research 2023, 29 (13), 2513–2524.

(35) Fauriat, C.; Long, E. O.; Ljunggren, H.-G.; Bryceson, Y. T. Regulation of human NK-cell cytokine and chemokine production by target cell recognition. Blood, The Journal of the American Society of Hematology 2010, 115 (11), 2167–2176.

(36) Öhlund, D.; Handly-Santana, A.; Biffi, G.; Elyada, E.; Almeida, A. S.; Ponz-Sarvise, M.; Corbo, V.; Oni, T. E.; Hearn, S. A.; Lee, E. J. Distinct populations of inflammatory fibroblasts and myofibroblasts in pancreatic cancer. Journal of Experimental Medicine 2017, 214 (3), 579–596.

(37) Chen, Y.; Kim, J.; Yang, S.; Wang, H.; Wu, C.-J.; Sugimoto, H.; LeBleu, V. S.; Kalluri, R. Type I collagen deletion in αSMA+ myofibroblasts augments immune suppression and accelerates progression of pancreatic cancer. Cancer cell 2021, 39 (4), 548–565. e546. DOI: 10.1016/j.ccell.2021.02.007.

(38) Lanier, L. L. NKG2D receptor and its ligands in host defense. Cancer immunology research 2015, 3 (6), 575–582.

(39) Carstens, J. L.; Correa de Sampaio, P.; Yang, D.; Barua, S.; Wang, H.; Rao, A.; Allison, J. P.; LeBleu, V. S.; Kalluri, R. Spatial computation of intratumoral T cells correlates with survival of patients with pancreatic cancer. Nat Commun 2017, 8, 15095. DOI: 10.1038/ncomms15095 From NLM Medline.

(40) Jin, S.; Guerrero-Juarez, C. F.; Zhang, L.; Chang, I.; Ramos, R.; Kuan, C.-H.; Myung, P.; Plikus, M. V.; Nie, Q. Inference and analysis of cell-cell communication using CellChat. Nature communications 2021, 12 (1), 1088. DOI: 10.1038/s41467-021-21246-9.

(41) Gulati, G. S.; Sikandar, S. S.; Wesche, D. J.; Manjunath, A.; Bharadwaj, A.; Berger, M. J.; Ilagan, F.; Kuo, A. H.; Hsieh, R. W.; Cai, S. Single-cell transcriptional diversity is a hallmark of developmental potential. Science 2020, 367 (6476), 405–411. DOI: 10.1126/science.aax0249.

(42) Enane, F. O.; Saunthararajah, Y.; Korc, M. Differentiation therapy and the mechanisms that terminate cancer cell proliferation without harming normal cells. Cell death & disease 2018, 9 (9), 912. DOI: 10.1038/s41419-018-0919-9.

(43) Zhang, T.; Ren, Y.; Yang, P.; Wang, J.; Zhou, H. Cancer-associated fibroblasts in pancreatic ductal adenocarcinoma. Cell Death Dis 2022, 13 (10), 897. DOI: 10.1038/s41419-022-05351-1 From NLM Medline.

(44) Wang, Y.; Liang, Y.; Xu, H.; Zhang, X.; Mao, T.; Cui, J.; Yao, J.; Wang, Y.; Jiao, F.; Xiao, X. Single-cell analysis of pancreatic ductal adenocarcinoma identifies a novel fibroblast subtype associated with poor prognosis but better immunotherapy response. Cell discovery 2021, 7 (1), 36.

(45) Clark, R. A.; Alon, R.; Springer, T. A. CD44 and hyaluronan-dependent rolling interactions of lymphocytes on tonsillar stroma. The Journal of cell biology 1996, 134 (4), 1075–1087.

(46) Hughes, C. S.; Postovit, L. M.; Lajoie, G. A. Matrigel: a complex protein mixture required for optimal growth of cell culture. Proteomics 2010, 10 (9), 1886–1890.

(47) Chen, C.; Zhao, S.; Karnad, A.; Freeman, J. W. The biology and role of CD44 in cancer progression: therapeutic implications. Journal of hematology & oncology 2018, 11, 1–23. DOI: 10.1186/s13045-018-0605-5.

(48) McMahon, M.; Ye, S.; Pedrina, J.; Dlugolenski, D.; Stambas, J. Extracellular matrix enzymes and immune cell biology. Frontiers in Molecular Biosciences 2021, 8, 703868. DOI: 10.3389/fmolb.2021.703868.

(49) Feig, C.; Jones, J. O.; Kraman, M.; Wells, R. J.; Deonarine, A.; Chan, D. S.; Connell, C. M.; Roberts, E. W.; Zhao, Q.; Caballero, O. L. Targeting CXCL12 from FAP-expressing carcinoma-associated fibroblasts synergizes with anti–PD-L1 immunotherapy in pancreatic cancer. Proceedings of the National Academy of Sciences 2013, 110 (50), 20212–20217.

(50) Paavola, K. J.; Roda, J. M.; Lin, V. Y.; Chen, P.; O’Hollaren, K. P.; Ventura, R.; Crawley, S. C.; Li, B.; Chen, H.-I. H.; Malmersjö, S. The fibronectin–ILT3 interaction functions as a stromal checkpoint that suppresses myeloid cells. Cancer immunology research 2021, 9 (11), 1283–1297.

(51) Özdemir, B. C.; Pentcheva-Hoang, T.; Carstens, J. L.; Zheng, X.; Wu, C.-C.; Simpson, T. R.; Laklai, H.; Sugimoto, H.; Kahlert, C.; Novitskiy, S. V. Depletion of carcinoma-associated fibroblasts and fibrosis induces immunosuppression and accelerates pancreas cancer with reduced survival. Cancer cell 2014, 25 (6), 719–734.

(52) Van Cutsem, E.; Tempero, M. A.; Sigal, D.; Oh, D.-Y.; Fazio, N.; Macarulla, T.; Hitre, E.; Hammel, P.; Hendifar, A. E.; Bates, S. E. Randomized phase III trial of pegvorhyaluronidase alfa with nab-paclitaxel plus gemcitabine for patients with hyaluronan-high metastatic pancreatic adenocarcinoma. Journal of Clinical Oncology 2020, 38 (27), 3185–3194.

(53) Delgado-Gonzalez, A.; Sanchez-Martin, R. M. Mass cytometry tags: where chemistry meets single-cell analysis. Analytical Chemistry 2020, 93 (2), 657–664. DOI: 10.1021/acs.analchem.0c03560.

(54) Windhager, J.; Zanotelli, V. R. T.; Schulz, D.; Meyer, L.; Daniel, M.; Bodenmiller, B.; Eling, N. An end-to-end workflow for multiplexed image processing and analysis. Nature Protocols 2023, 1–49. DOI: 10.1038/s41596-023-00881-0.

(55) Lee, C.-W.; Ren, Y. J.; Marella, M.; Wang, M.; Hartke, J.; Couto, S. S. Multiplex immunofluorescence staining and image analysis assay for diffuse large B cell lymphoma. Journal of immunological methods 2020, 478, 112714. DOI: 10.1016/j.jim.2019.112714.

(56) Bankhead, P.; Loughrey, M. B.; Fernández, J. A.; Dombrowski, Y.; McArt, D. G.; Dunne, P. D.; McQuaid, S.; Gray, R. T.; Murray, L. J.; Coleman, H. G. QuPath: Open source software for digital pathology image analysis. Scientific reports 2017, 7 (1), 1–7.

(57) Schmidt, U.; Weigert, M.; Broaddus, C.; Myers, G. Cell detection with star-convex polygons. In International conference on medical image computing and computer-assisted intervention, 2018; Springer: pp 265–273.

(58) Scrucca, L.; Fop, M.; Murphy, T. B.; Raftery, A. E. mclust 5: clustering, classification and density estimation using Gaussian finite mixture models. The R journal 2016, 8 (1), 289. DOI: 10.32614/RJ-2016-021.

(59) Zahavi, D. J.; Erbe, R.; Zhang, Y.-W.; Guo, T.; Malchiodi, Z. X.; Maynard, R.; Lekan, A.; Gallagher, R.; Wulfkuhle, J.; Petricoin, E. Antibody dependent cell-mediated cytotoxicity selection pressure induces diverse mechanisms of resistance. Cancer Biology & Therapy 2023, 24 (1), 2269637. DOI: 10.1080/15384047.2023.2269637.

(60) Ajina, R.; Malchiodi, Z. X.; Fitzgerald, A. A.; Zuo, A.; Wang, S.; Moussa, M.; Cooper, C. J.; Shen, Y.; Johnson, Q. R.; Parks, J. M. Antitumor T-cell immunity contributes to pancreatic cancer immune resistance. Cancer immunology research 2021, 9 (4), 386–400.

(61) Carr, W. H.; Zinsou, C.; Rowe, T.; Lange, C.; Lawal, O. O. Human natural killer cell expansion in vitro using mitomycin-C treatment as an alternative to irradiation of modified K562 feeder cells. The Journal of Immunology 2017, 198 (1_Supplement), 82.16–82.16. DOI: 10.4049/jimmunol.198.Supp.82.16.

(62) Schindelin, J.; Arganda-Carreras, I.; Frise, E.; Kaynig, V.; Longair, M.; Pietzsch, T.; Preibisch, S.; Rueden, C.; Saalfeld, S.; Schmid, B. Fiji: an open-source platform for biological-image analysis. Nature methods 2012, 9 (7), 676–682. DOI: 10.1038/nmeth.2019.

