## Supplementary Materials for "Periductal Fibroblast Density Defines Lymphocyte Exclusion via a CD44-Dependent Stromal Checkpoint in Pancreatic Cancer"

### Supplementary Figures

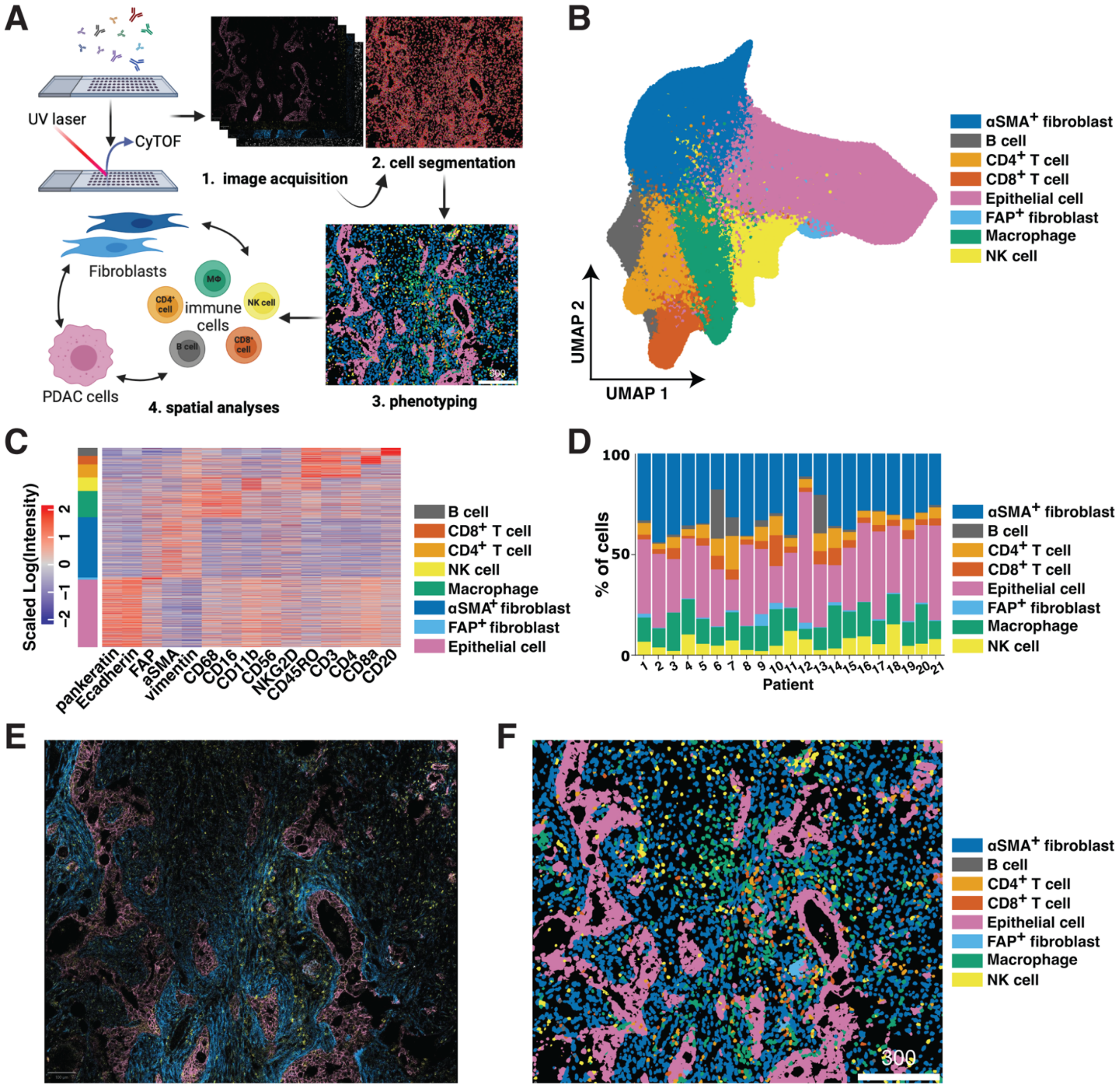


**Supplementary Figure S1: Spatial proteomic analysis of the human PDAC TME resolves PDAC cell populations. A.** Schematic of IMC workflow. **B.** UMAP of IMC-defined cell populations from 21 untreated PDAC patient samples. **C.** Heatmap of scaled log-normalized protein expression intensities of canonical marker expression used to phenotype segmented human PDAC cell populations. 15k cells randomly subsampled from the 21 PDAC patient samples are displayed. **D.** Bar plot showing percentages of cell populations per PDAC patient. **E.** Representative IMC pseudoimage of the human PDAC TME highlighting distinctions between epithelial cells (Pan-cytokeratin and E-cadherin; pink), fibroblasts (αSMA and FAP; blue), and immune cells (CD45RO; yellow). **F.** Single cell image (Cytomapper) of **E** displaying all identified cell types from **B**-**D**.

**A**

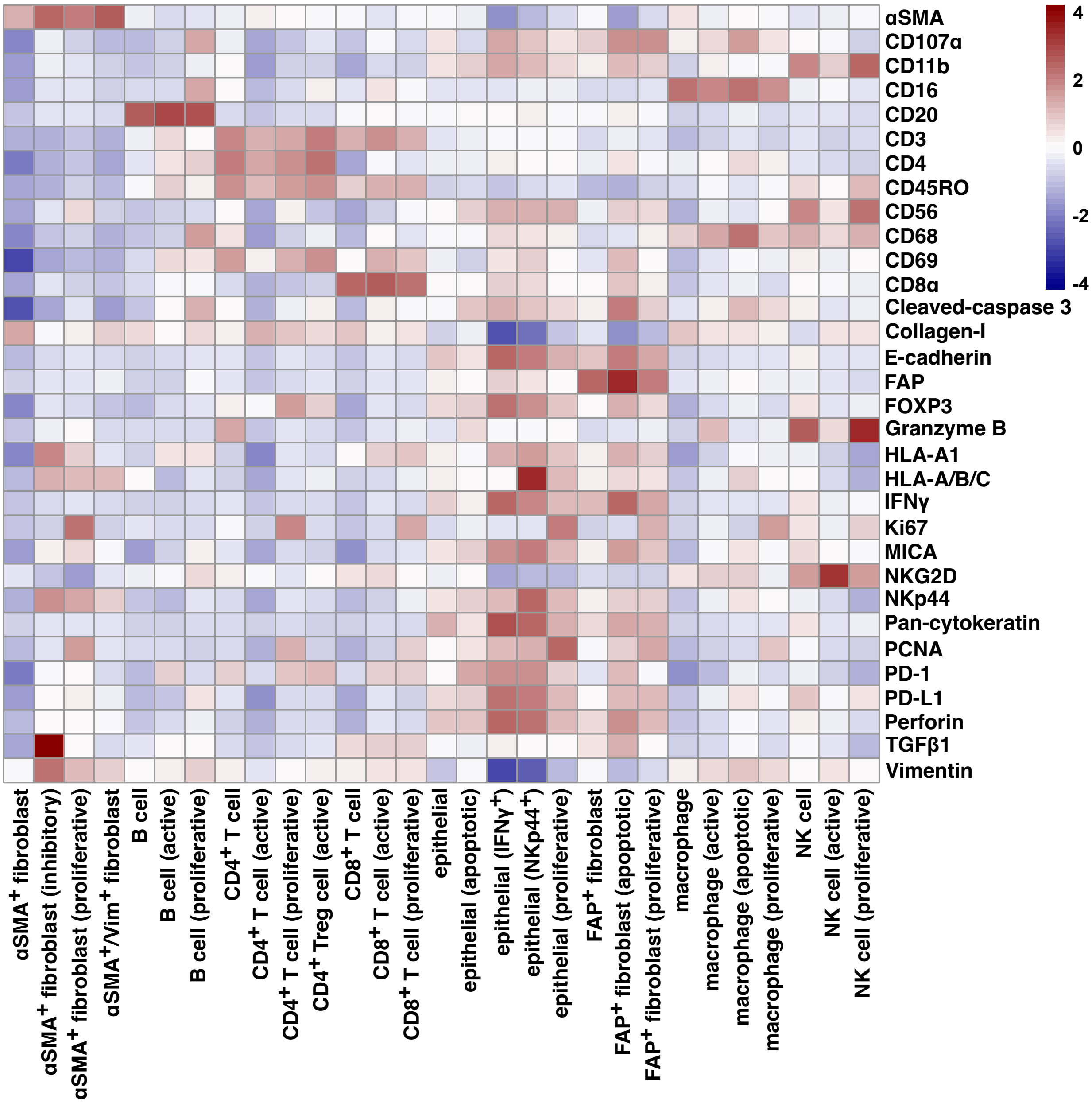

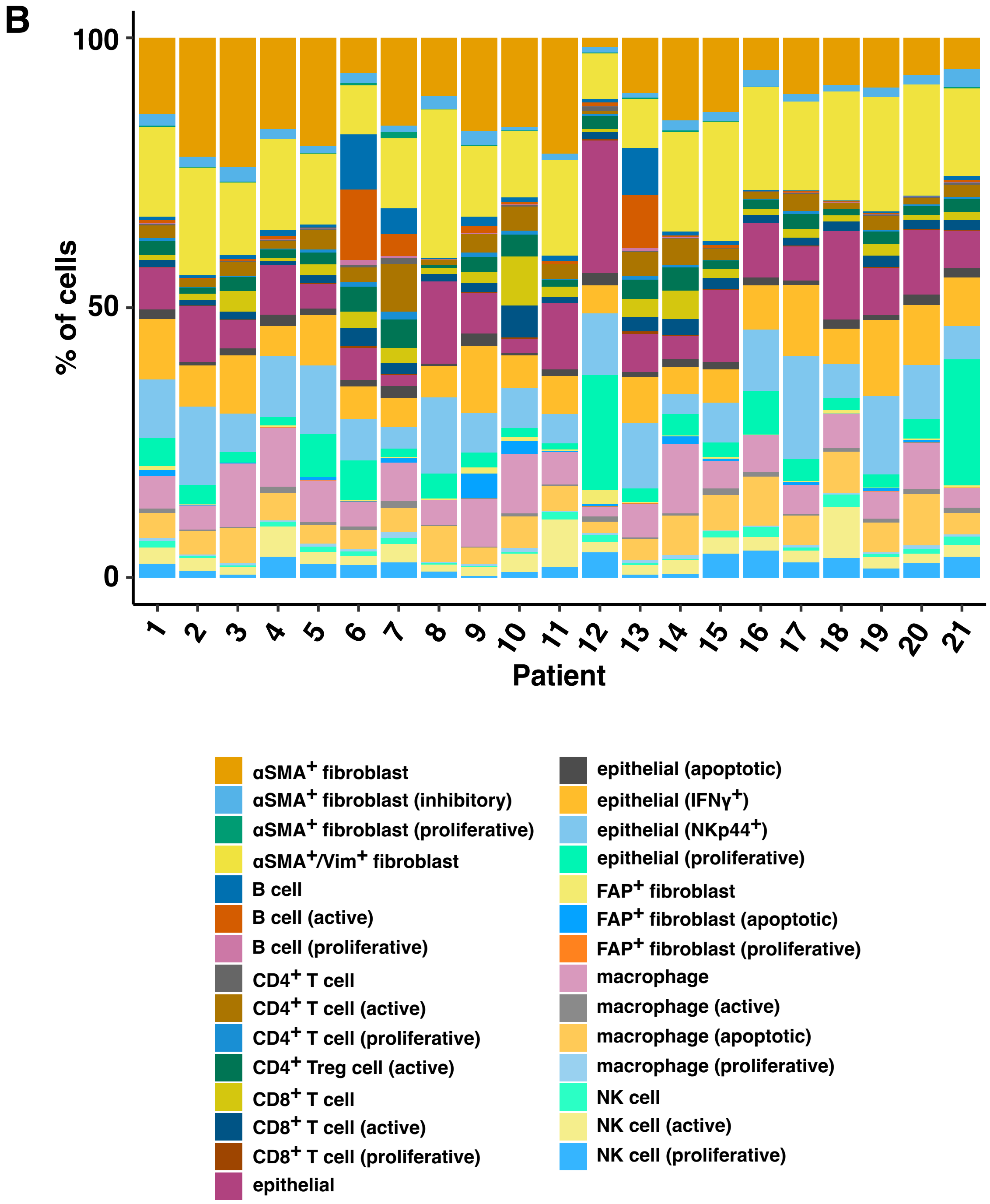


**Supplementary Figure S2: Subclustered cell populations of spatial proteomic resolved PDAC cell populations. A.** Averaged expression heatmap of IMC-identified PDAC subclustered cell populations. **B.** Bar plot showing percentages of subclustered cell populations per PDAC patient.

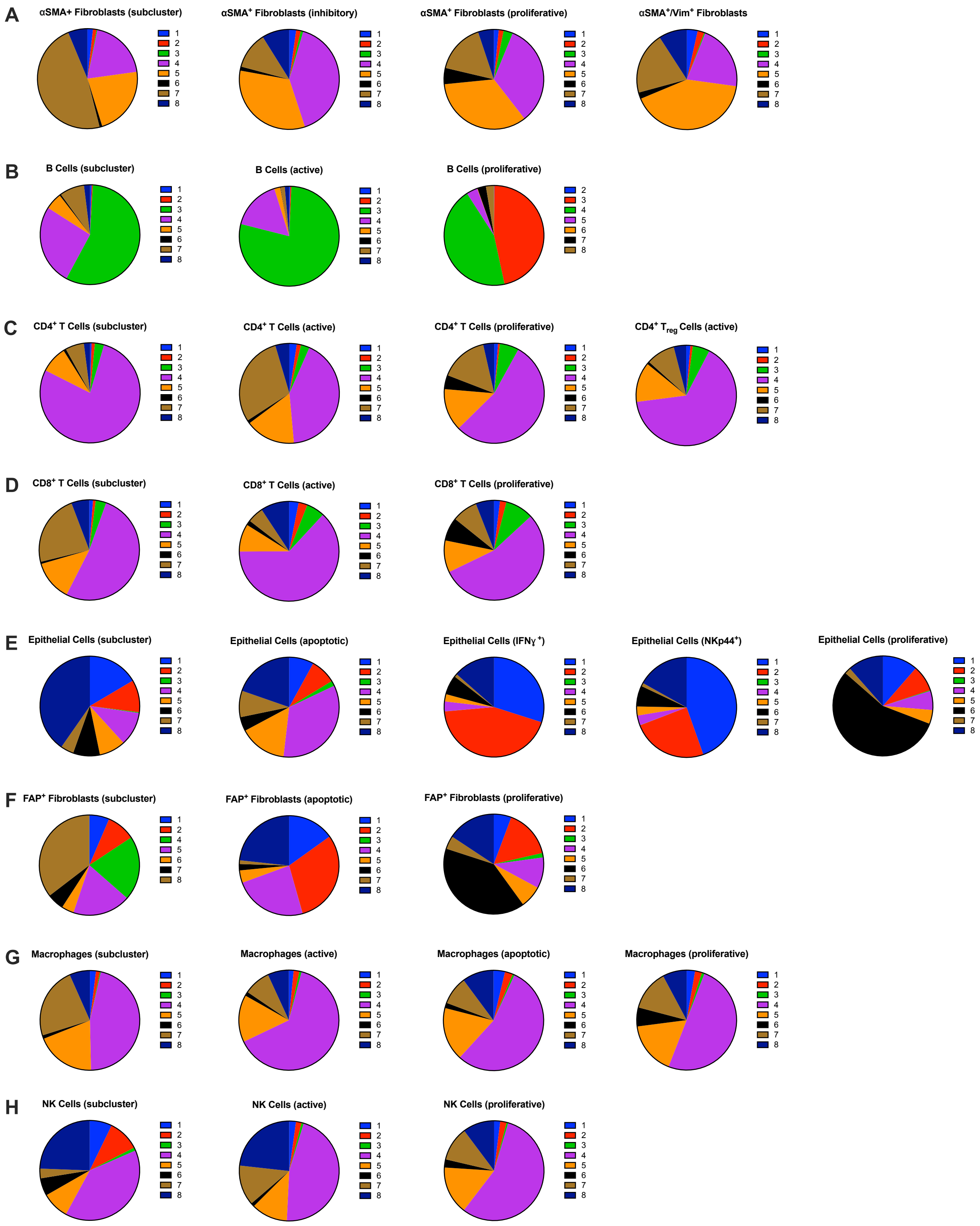


**Supplementary Figure S3: Proportions of PDAC subclustered cell populations per cellular neighborhood.** Pie charts depicting proportions of **A.** αSMA$^{+}$ fibroblast subclusters (left to right:

αSMA$^+$ fibroblast (subcluster), αSMA$^+$ fibroblast (inhibitory), αSMA$^+$ fibroblast (proliferative), αSMA$^+$/Vim$^+$ fibroblast), **B.** B cell subclusters (left to right: B cell (subcluster), B cell (active), B cell (proliferative))m, **C.** CD4$^+$ T cell subclusters (left to right: CD4$^+$ T cell (subcluster), CD4$^+$ T cell (active), CD4$^+$ T cell (proliferative), CD4$^+$ Treg cell (active)), **D.** CD8$^+$ T cell subclusters (left to right: CD8$^+$ T cell (subcluster), CD8$^+$ T cell (active), CD8$^+$ T cell (proliferative)), **E.** Epithelial cell subclusters (left to right: epithelial (subcluster), epithelial (apoptotic), epithelial (IFNγ$^+$), epithelial (NKp44$^+$), epithelial (proliferative)), **F.** FAP$^+$ fibroblast subclusters (left to right: FAP$^+$ fibroblast (subcluster), FAP$^+$ fibroblast (apoptotic), FAP$^+$ fibroblast (proliferative)), **G.** Macrophage subclusters (left to right: macrophage (subcluster), macrophage (active), macrophage (apoptotic), macrophage (proliferative)), and **H.** NK cell subclusters (left to right: NK cell (subcluster), NK cell (active), NK cell (proliferative)) per cellular neighborhood.

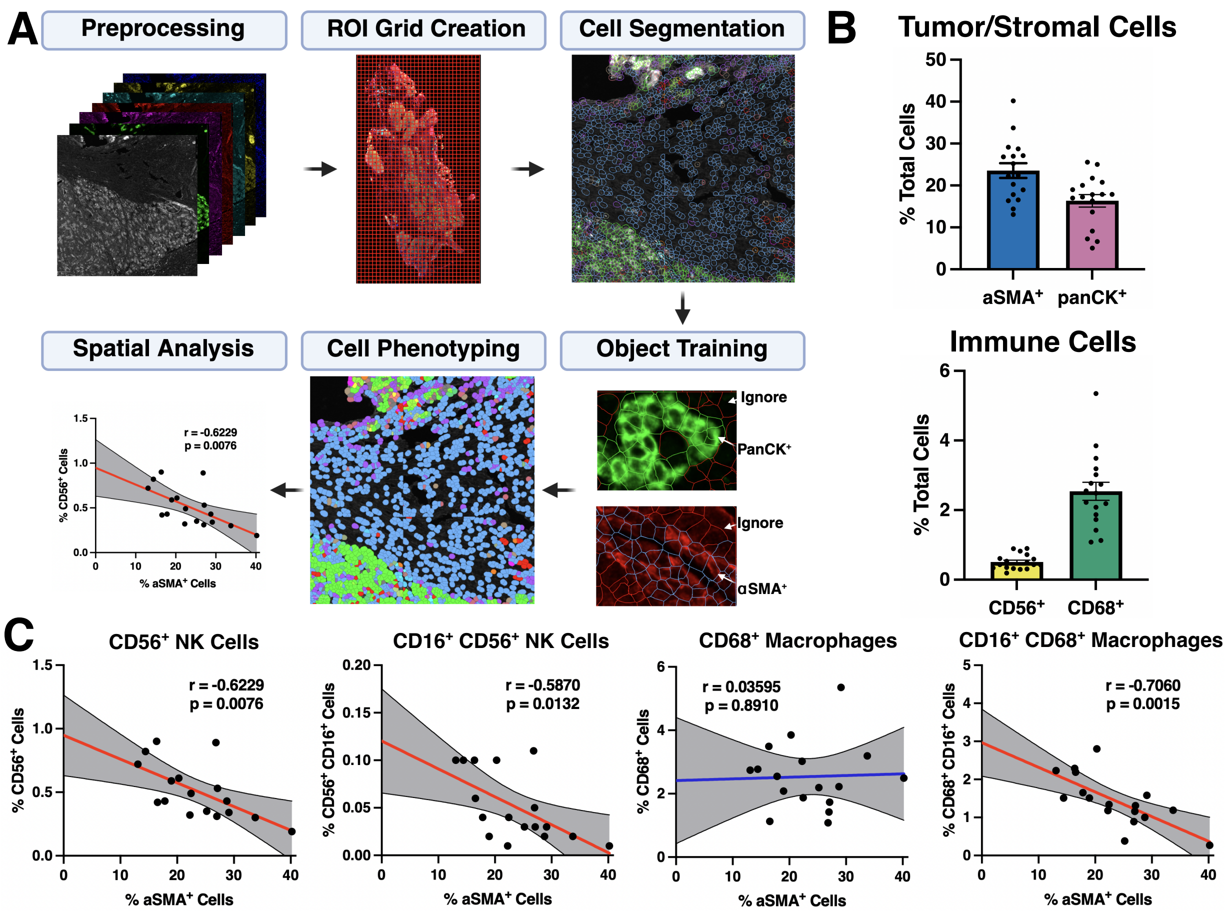


**Supplementary Figure S4: Pro-inflammatory NK cell and Macrophage content is inversely correlated with αSMA$^+$ fibroblasts. A.** Schematic of mfIHC workflow. **B.** Barplot showing representative cell populations per PDAC sample (n=17). **C.** Correlations between specific leukocyte populations and αSMA+ fibroblasts with Pearson's correlation. Gray bands represent 95% confidence bands of the best-fit line.

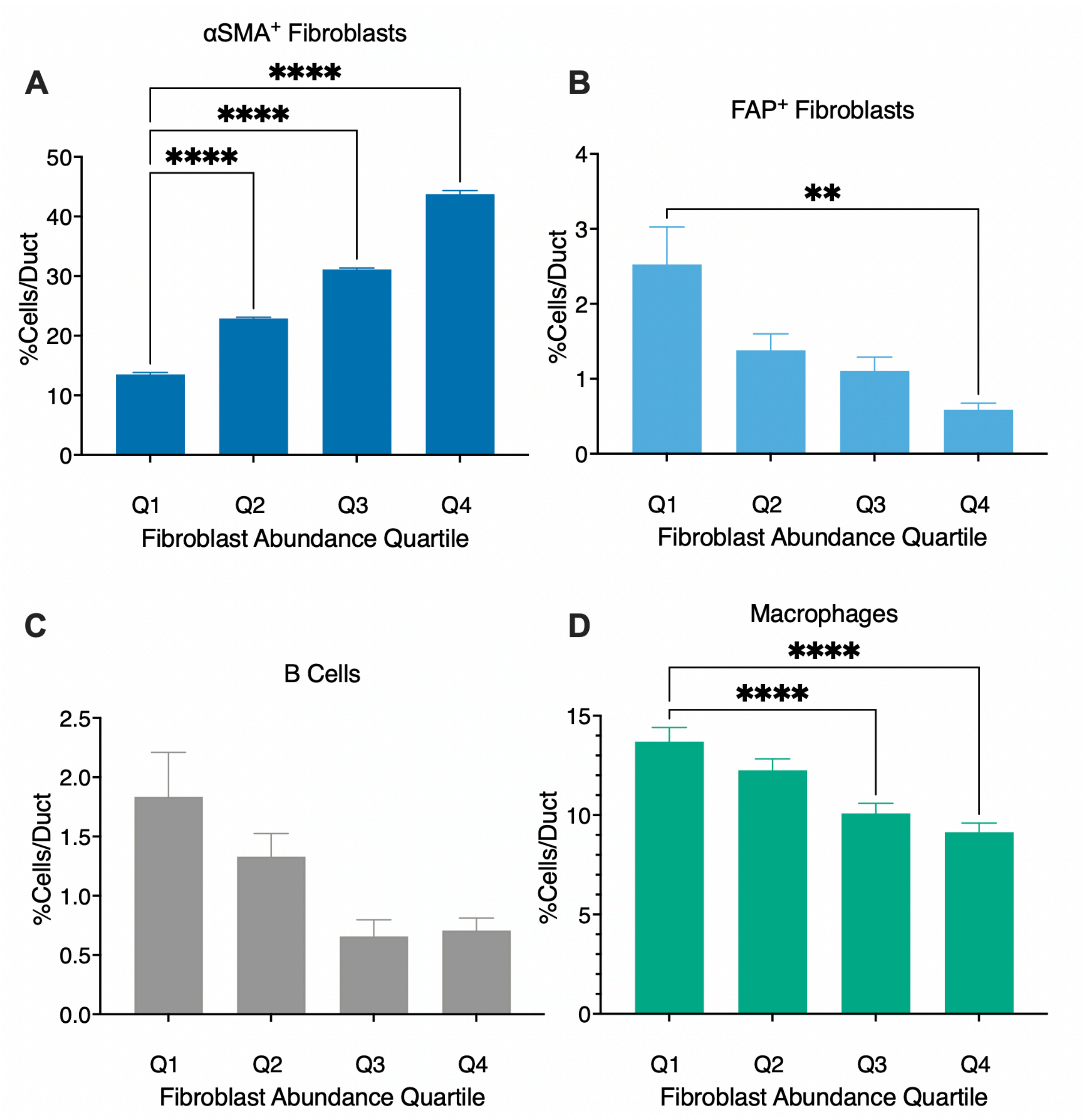


**Supplementary Figure S5: PDAC epithelial-ductal ROI cell type abundance.** Abundance of **A.** αSMA+ fibroblasts, **B.** FAP+ fibroblasts, **C.** B Cells and **D.** macrophages in epithelial-ductal ROIs ranked by fibroblast abundance quartile (Q1-Q4) represented as % cells/duct. 2-Way ANOVA (*$p < 0.05$, **$p < 0.01$, ***$p < 0.001$, ****$p < 0.0001$; n = 952, total epithelial-ductal ROIs).

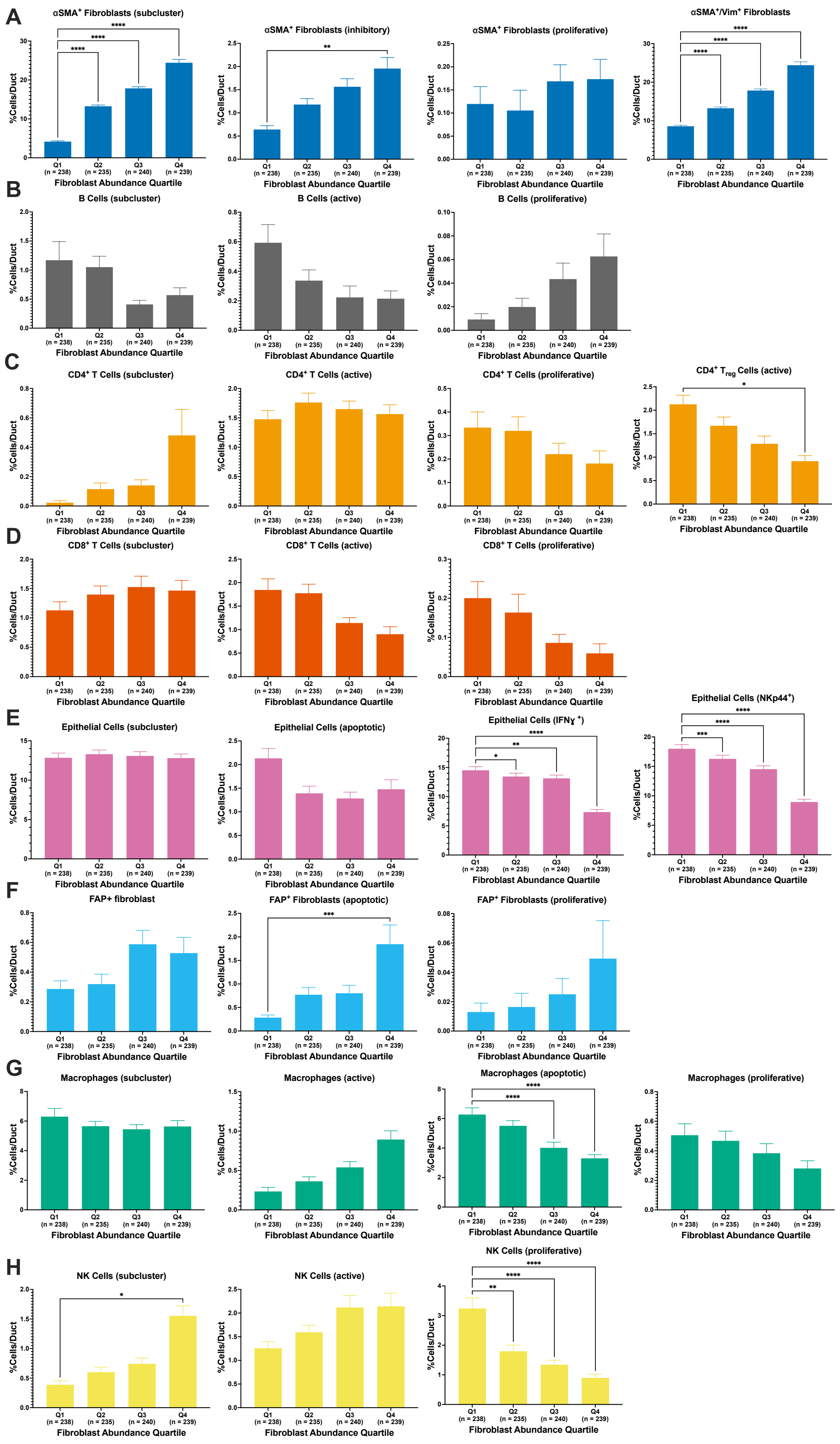
A
αSMA+ Fibroblasts (subcluster)
αSMA+ Fibroblasts (inhibitory)
αSMA+ Fibroblasts (proliferative)
αSMA+/Vim+ Fibroblasts
B
B Cells (subcluster)
B Cells (active)
B Cells (proliferative)
C
CD4+ T Cells (subcluster)
CD4+ T Cells (active)
CD4+ T Cells (proliferative)
CD4+ Treg Cells (active)
D
CD8+ T Cells (subcluster)
CD8+ T Cells (active)
CD8+ T Cells (proliferative)
E
Epithelial Cells (subcluster)
Epithelial Cells (apoptotic)
Epithelial Cells (IFNɣ +)
Epithelial Cells (NKp44+)
F
FAP+ fibroblast
FAP+ Fibroblasts (apoptotic)
FAP+ Fibroblasts (proliferative)
G
Macrophages (subcluster)
Macrophages (active)
Macrophages (apoptotic)
Macrophages (proliferative)
H
NK Cells (subcluster)
NK Cells (active)
NK Cells (proliferative)
%Cells/Duct
Q1 (n = 238)
Q2 (n = 235)
Q3 (n = 240)
Q4 (n = 239)
Fibroblast Abundance Quartile

52

**Supplementary Figure S6: PDAC epithelial-ductal ROI cell subcluster abundance.** Abundance of **A.** αSMA$^{+}$ fibroblast subclusters (left to right: αSMA$^{+}$ fibroblast (subcluster), αSMA$^{+}$ fibroblast (inhibitory), αSMA$^{+}$ fibroblast (proliferative), αSMA$^{+}$/Vim$^{+}$ fibroblast), **B.** B cell subclusters (left to right: B cell (subcluster), B cell (active), B cell (proliferative)), **C.** CD4$^{+}$ T cell subclusters (left to right: CD4$^{+}$ T cell (subcluster), CD4$^{+}$ T cell (active), CD4$^{+}$ T cell (proliferative), CD4$^{+}$ Treg cell (active)), **D.** CD8$^{+}$ T cell subclusters (left to right: CD8$^{+}$ T cell (subcluster), CD8$^{+}$ T cell (active), CD8$^{+}$ T cell (proliferative)), **E.** epithelial cell subclusters (left to right: epithelial (subcluster), epithelial (apoptotic), epithelial (IFNγ$^{+}$), epithelial (NKp44$^{+}$), **F.** FAP$^{+}$ fibroblast subclusters (left to right: FAP$^{+}$ fibroblast (subcluster), FAP$^{+}$ fibroblast (apoptotic), FAP$^{+}$ fibroblast (proliferative)), **G.** macrophage subclusters (left to right: macrophage (subcluster), macrophage (active), macrophage (apoptotic), macrophage (proliferative)), and **H.** NK cell subclusters (left to right: NK cell (subcluster), NK cell (active), NK cell (proliferative)) in epithelial-ductal ROIs ranked by fibroblast abundance quartile (Q1-A4) represented as % cells/duct. 2-Way ANOVA (*$p < 0.05$, **$p < 0.01$, ***$p < 0.001$, ****$p < 0.0001$; n = 952, total epithelial-ductal ROIs).

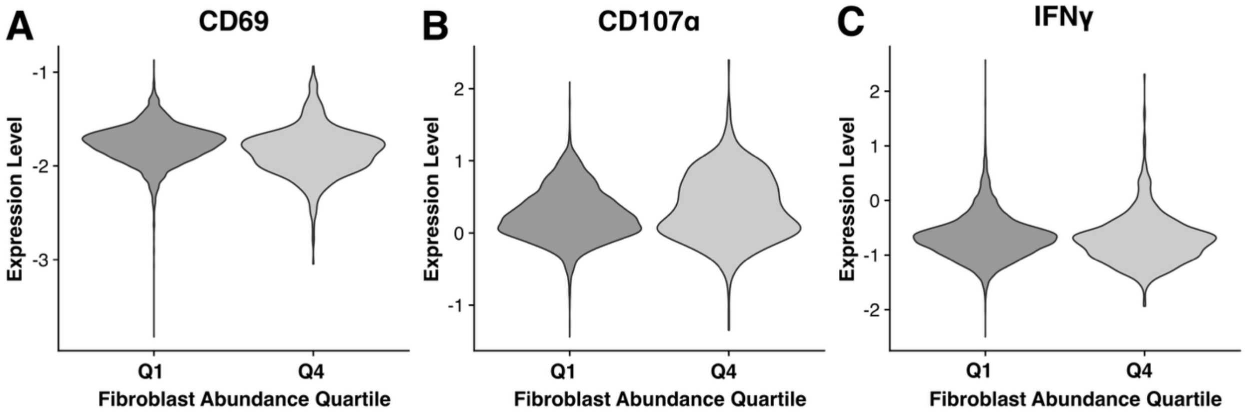


**Supplementary Figure S7: NK cells retain expression of activator markers in fibroblast-rich periductal regions.** Violin plot showing NK cell-based expression in Q1 and Q4 fibroblast abundance quartiles of **A.** CD69, **B.** CD107a, and **C.** IFNγ (not significant, Kruskal-Wallis test).

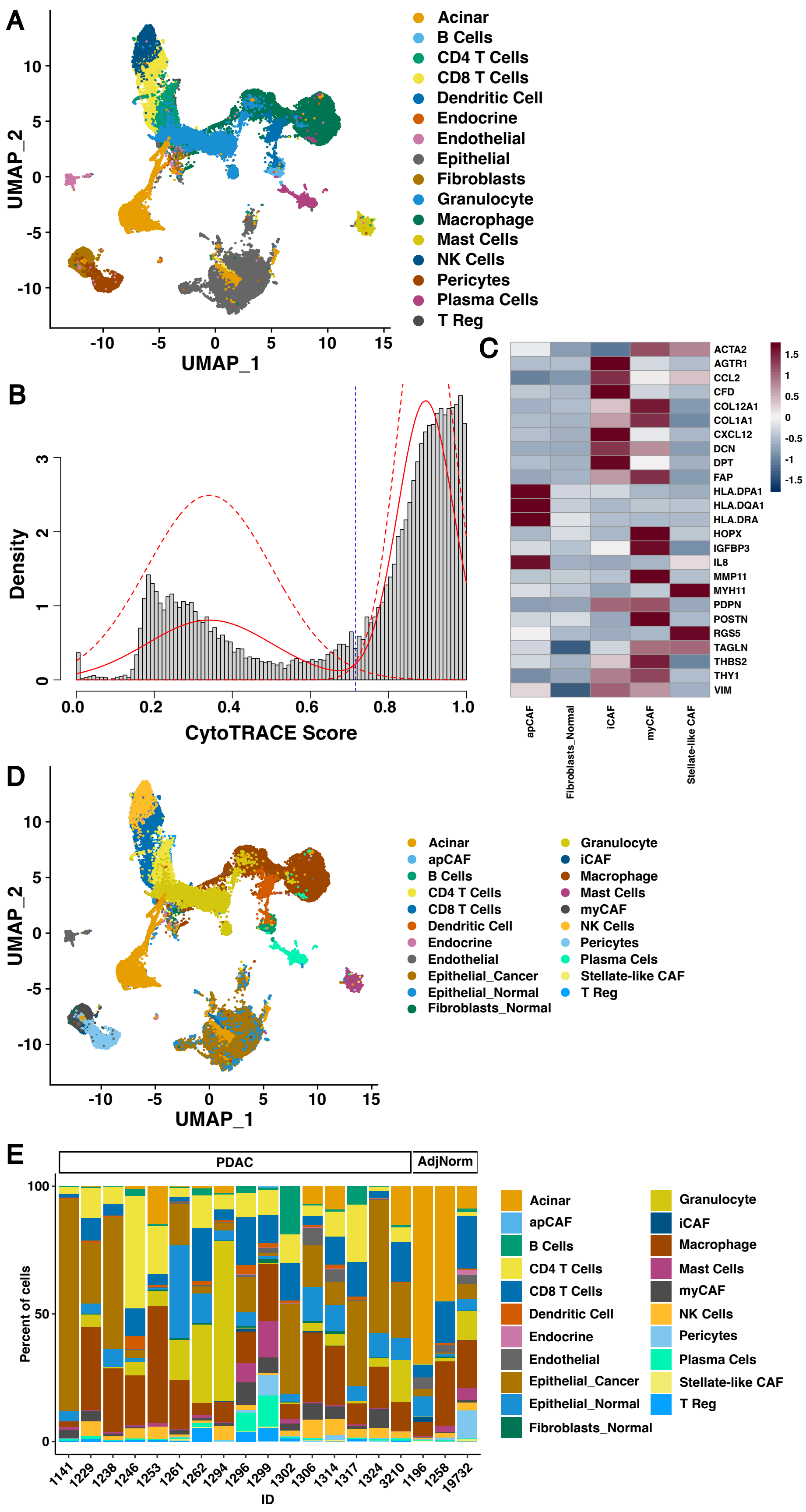
A
UMAP_2
UMAP_1
Acinar
B Cells
CD4 T Cells
CD8 T Cells
Dendritic Cell
Endocrine
Endothelial
Epithelial
Fibroblasts
Granulocyte
Macrophage
Mast Cells
NK Cells
Pericytes
Plasma Cells
T Reg
B
Density
CytoTRACE Score
C
ACTA2
AGTR1
CCL2
CFD
COL12A1
COL1A1
CXCL12
DCN
DPT
FAP
HLA.DPA1
HLA.DQA1
HLA.DRA
HOPX
IGFBP3
IL8
MMP11
MYH11
PDPN
POSTN
RGS5
TAGLN
THBS2
THY1
VIM
apCAF
Fibroblasts_Normal
iCAF
myCAF
Stellate-like CAF
D
UMAP_2
UMAP_1
Acinar
apCAF
B Cells
CD4 T Cells
CD8 T Cells
Dendritic Cell
Endocrine
Endothelial
Epithelial_Cancer
Epithelial_Normal
Fibroblasts_Normal
Granulocyte
iCAF
Macrophage
Mast Cells
myCAF
NK Cells
Pericytes
Plasma Cels
Stellate-like CAF
T Reg
E
PDAC
AdjNorm
Percent of cells
ID
1141
1229
1238
1246
1253
1261
1262
1294
1296
1299
1302
1306
1314
1317
1324
3210
1196
1258
19732
Acinar
apCAF
B Cells
CD4 T Cells
CD8 T Cells
Dendritic Cell
Endocrine
Endothelial
Epithelial_Cancer
Epithelial_Normal
Fibroblasts_Normal
Granulocyte
iCAF
Macrophage
Mast Cells
myCAF
NK Cells
Pericytes
Plasma Cels
Stellate-like CAF
T Reg

**Supplementary Figure S8: Subclustering of epithelial and fibroblast populations in human PDAC scRNAseq samples. A.** UMAP of all PDAC and AdjNorm tissue from Steele et al. cell annotations. **B.** Histogram of CytoTRACE scores from subsetted, re-integrated epithelial cells displaying two distinct epithelial cell populations with cutoff at 0.72 (blue line) for downstream analyses. **C.** Heatmap of subclustered fibroblasts populations with known fibroblasts population genes. **D.** UMAP of relabeled cell populations after subclustering of epithelial cell and fibroblast subpopulations colored from original annotations distinguishing normal and malignant epithelial cells, iCAFs, myCAFs, apCAFs, normal fibroblasts, and stellate-like CAFs. **E.** Percentage of relabeled cells per patient in PDAC and AdjNorm tissue.

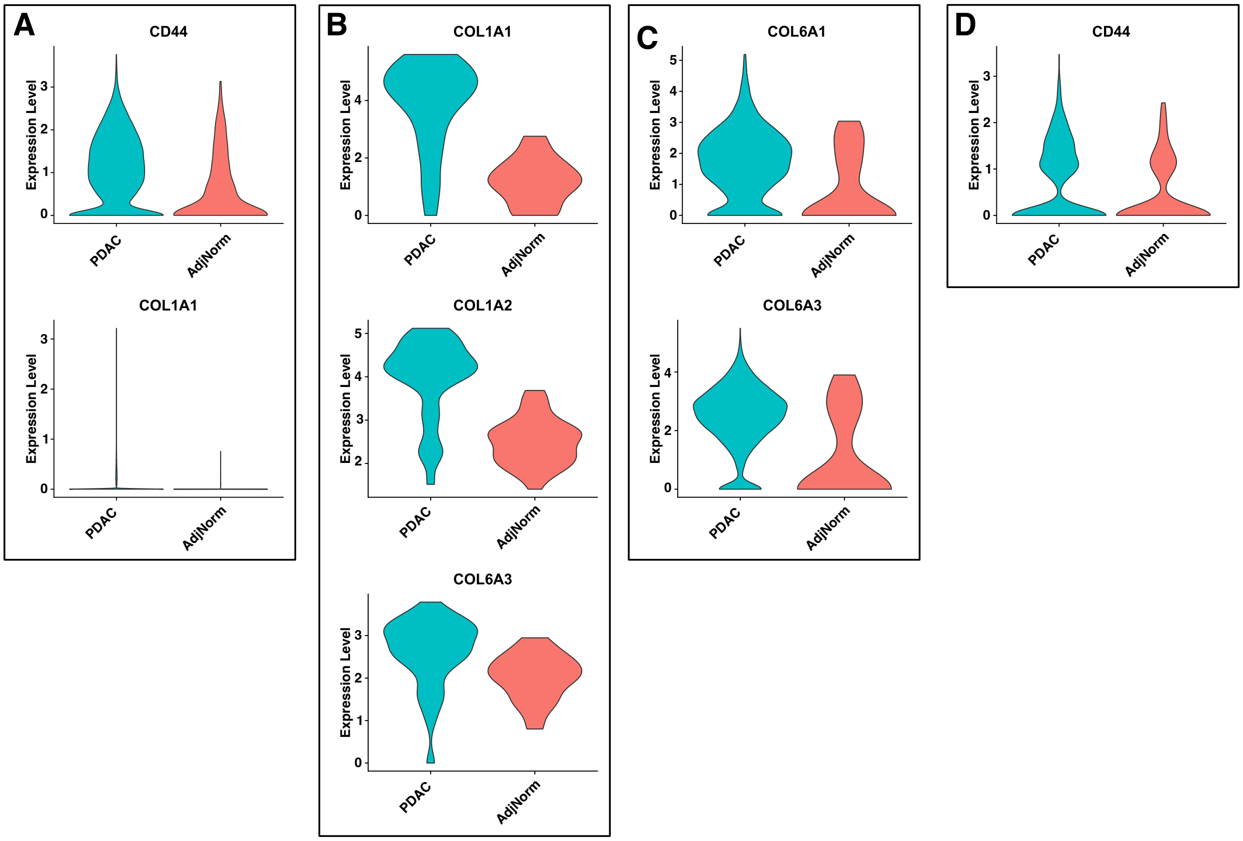


**Supplementary Figure S9: Fibroblast and epithelial cell populations exhibit differential expression of CellChat-based collagen signaling genes.** Violin plots showing differential expression of genes associated with CellChat-identified ligand-receptor pairs between PDAC (teal) and AdjNorm (pink) in **A.** malignant epithelial cells, **B.** iCAFs, and **C.** myCAFs, and **D.** NK cells (adjusted-$P < 0.01$, Bonferroni correction).

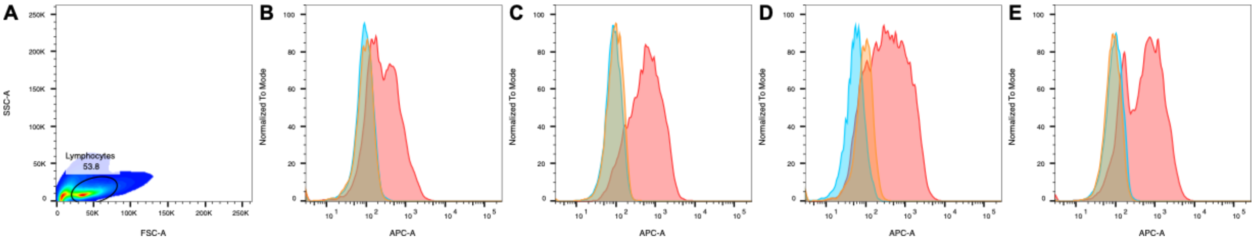


**Supplementary Figure S10: Human donor NK cells express CD44. A.** Representative gating of donor NK cell lines. Flow cytometry histograms for CD44 expression on **B.** Donor NK #1, **C.** Donor NK #2, **D.** Donor NK #3, and **E.** Donor NK #4 (orange = unstained, blue = Fc block only control, red = Fc block + α-CD44).

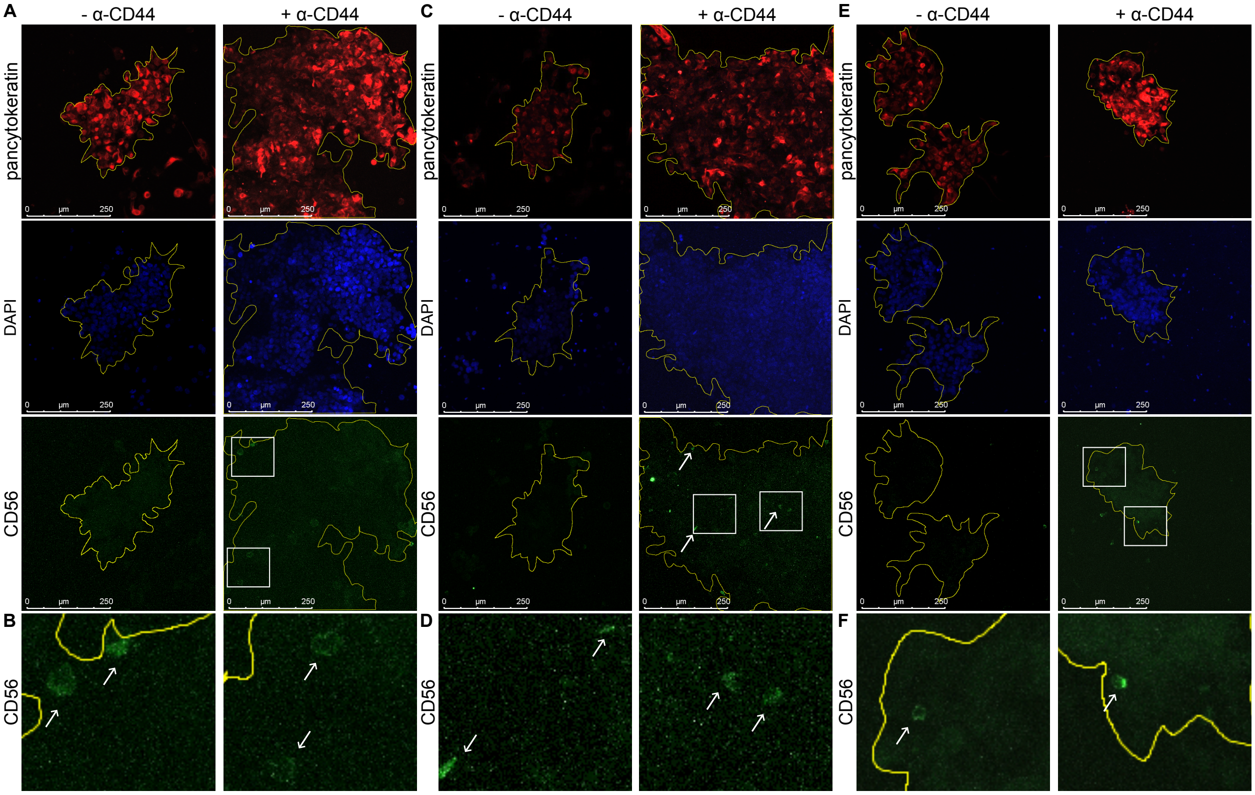

**Supplementary Figure S11: CD44 neutralization increases human donor NK cell invasion into PANC-1 PDAC spheroids.** Representative 20X immunofluorescence images of outlined (yellow) PANC-1 spheroids (Pan-cytokeratin; red) embedded with **A.** Donor NK #1, **C.** Donor NK #2, and **E.** Donor NK #3 cell lines (CD56; green) treated with or without α-CD44 (DAPI = blue). Scale bar = 250 μm. Zoomed inset of **B.** Donor NK #1, **D.** Donor NK #2, and **F.** Donor NK #3 cell lines in the α-CD44 treatment group from **A**, **C**, and **E** (white boxes).

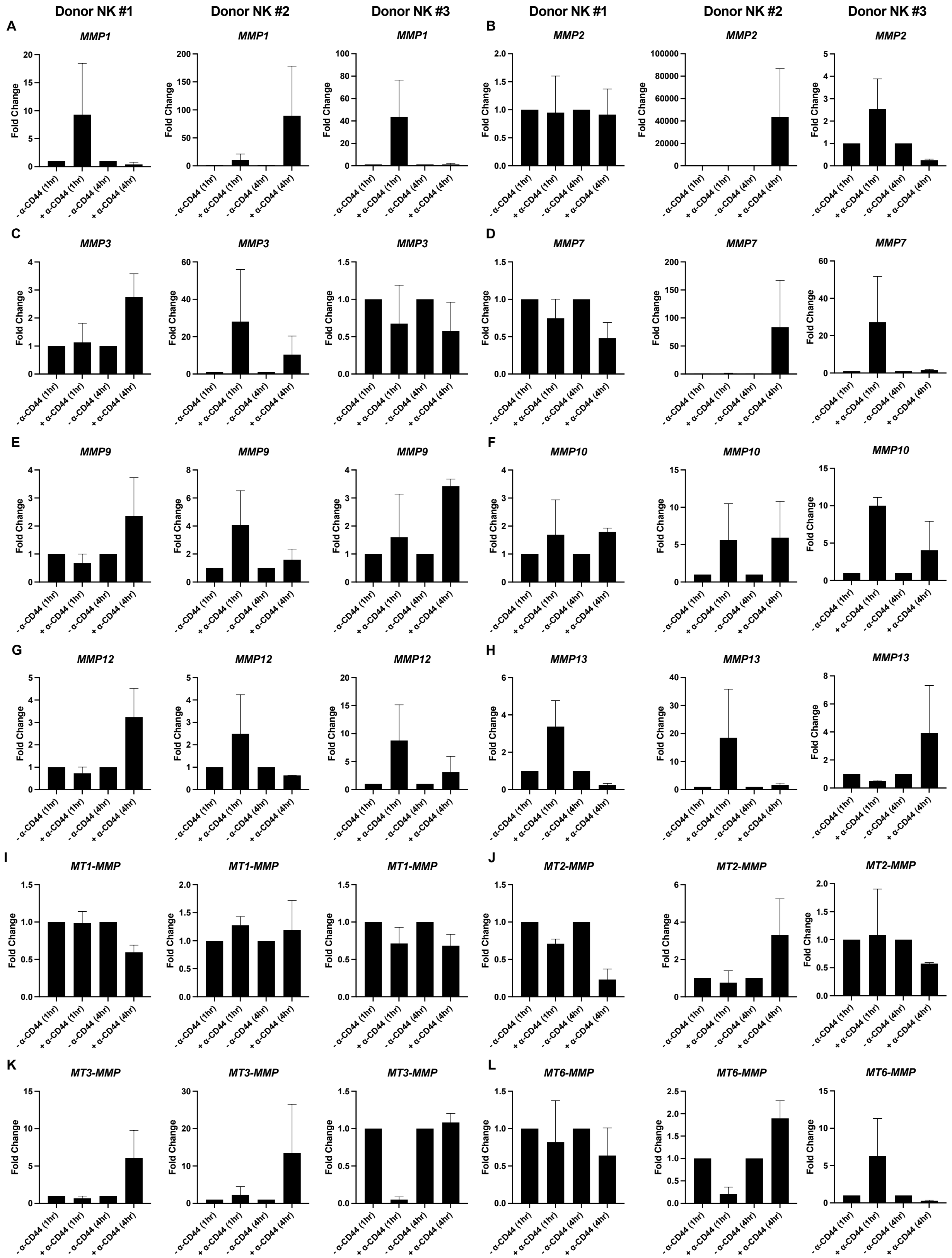
Donor NK #1
Donor NK #2
Donor NK #3
Donor NK #1
Donor NK #2
Donor NK #3
A
MMP1
B
MMP2
C
MMP3
D
MMP7
E
MMP9
F
MMP10
G
MMP12
H
MMP13
I
MT1-MMP
J
MT2-MMP
K
MT3-MMP
L
MT6-MMP
Fold Change
- α-CD44 (1hr)
+ α-CD44 (1hr)
- α-CD44 (4hr)
+ α-CD44 (4hr)

**Supplementary Figure S12: CD44 neutralization does not affect expression of MMP genes in human donor NK cells.** Bar plots of mean fold change ± SEM of the following MMP genes in three human donor NK cell lines: **A.** *MMP1*, **B.** *MMP2*, **C.** *MMP3*, **D.** *MMP7*, **E.** *MMP9*, **F.** *MMP10*, **G.** *MMP12*, **H.** *MMP13*, **I.** *MT1-MMP*, **J.** *MT2-MMP*, **K.** *MT3-MMP*, **L.** *MT6-MMP* (n = 2/line; Šídák's multiple comparisons test).

#### Supplementary Tables

**Supplementary Table S1: IMC antibody panel and metal assignments.**

**Maxpar® Human Immuno-Oncology IMC Panel Kit**

| Target | Company | Catalog Number | Metal Tag | Dilution |
|---|---|---|---|---|
| CD20 | Standard BioTools Inc. (SBI) | 201508 | 161Dy | 1:50 |
| CD3 | | | 170Er | 1:50 |
| CD4 | | | 156Gd | 1:50 |
| CD45RO | | | 173Yb | 1:50 |
| CD68 | | | 159Tb | 1:50 |
| CD8a | | | 162Dy | 1:50 |
| FoxP3 | | | 155Gd | 1:50 |
| Pan-keratin | | | 148Nd | 1:150 |
| Granzyme B | | | 167Er | 1:50 |
| Ki-67 | | | 168Er | 1:50 |
| PD-1 | | | 165Ho | 1:50 |
| PD-L1 | | | 150Nd | 1:50 |
| αSMA | | | 141Pr | 1:150 |
| Collagen-I | | | 169Tm | 1:50 |
| E-cadherin | | | 158Gd | 1:50 |
| Vimentin | | | 143Nd | 1:450 |
| Nucleic acid | | | 191Ir/193Ir | 1:50 |

**Maxpar® X8 Antibodies**

| Target | Company | Catalog Number | Metal Tag | Dilution |
|---|---|---|---|---|
| Perforin | SBI | 3176026D | 176Yb | 1:20 |
| Cleaved caspase-3 | | 3172027D | 172Yb | 1:50 |
| HLA-ABC | | 3144027D | 144Nd | 1:20 |
| CD107α (LAMP1) | | 3151021D | 151Eu | 1:150 |
| CD16 | | 3146020D | 146Nd | 1:50 |
| CD11b | | 3149028D | 149Sm | 1:50 |

**Non-Maxpar® Antibodies and Maxpar® X8 Antibody Labeling Kits**

| Target | Company | Product # | Dilution | Maxpar® X8 Antibody Labeling Kits | Catalog Number |
|---|---|---|---|---|---|
| Fibroblast Activation Protein (FAP) | Abcam | ab271976 | 1:50 | 153Eu | 201153A |
| CD56 | BioLegend | 304602 | 1:20 | 166Er | 201166A |
| TGFβ1 | Santa Cruz Biotechnology | sc-52893 | 1:750 | 163Dy | 201163A |

| **Supplementary Table S1 (continued)** | | | | | |
|---|---|---|---|---|---|
| Interferon (IFN)-γ | Abcam | ab218890 | 1:50 | 160Gd | 201160A |
| MICA | BioLegend | 320902 | 1:25 | 152Sm | 201152A |
| PCNA | Abcam | ab264494 | 1:50 | 147Sm | 201147A |
| NKG2D (CD314) | Novus Biologicals | NB100-65956 | 1:50 | 175Lu | 201175A |
| NKp44 | BioLegend | 325102 | 1:20 | 145Nd | 201145A |
| CD69 | Abcam | ab234512 | 1:20 | 174Yb | 201174A |
| HLA-A1 | Abcam | ab216653 | 1:50 | 164Dy | 201164A |

**Supplementary Table S2: PDAC patient information for IMC.**

| Patient | Case Number | Tumor size (cm) | Age (y) | Stage | Ethnicity | Race | Sex |
|---|---|---|---|---|---|---|---|
| 1 | 1A0021 | 2.9 | 78 | T2 N1 Mx | non-hispanic | White | F |
| 2 | 1A0043 | 3.5 | 69 | T3 N0 Mx | non-hispanic | Asian | M |
| 3 | 1A0122 | 3 | 67 | T3 N1 Mx | non-hispanic | African American | F |
| 4 | 1A0141 | 2 | 64 | T3 N1 Mx | non-hispanic | White | F |
| 5 | 1A0251 | 3.4 | 65 | T2 N0 | non-hispanic | African American | M |
| 6 | 1A0264 | 4.1 | 53 | T3 N1 M0 | non-hispanic | White | M |
| 7 | 1A0339 | 4.8 | 70 | T2 N1 | non-hispanic | African American | F |
| 8 | 1A0423 | 3 | 60 | T2 N1 | non-hispanic | White | F |
| 9 | 1A0445 | 3.5 | 68 | T3 N0 | non-hispanic | African American | F |
| 10 | 1A0459 | 3.8 | 60 | T2 N1 | non-hispanic | White | F |
| 11 | 1A0475 | 5 | 65 | T3 N1 M1 | non-hispanic | White | F |
| 12 | 1A0651 | 2.2 | 74 | T3 N1 | non-hispanic | African American | F |
| 13 | 1A0652 | 1.7 | 65 | T1 pN1 | non-hispanic | unknown | M |
| 14 | 1A0729 | 4.5 | 41 | T3 N1 | non-hispanic | White | F |
| 15 | 1A0803 | 4.3 | 63 | T3 N1 | non-hispanic | White | M |
| 16 | 1A0904 | 2 | 78 | T3 N1 | non-hispanic | White | F |
| 17 | 1A1105 | 2.3 | 64 | T3 N0 | non-hispanic | White | M |
| 18 | 1A1128 | 2.2 | 57 | T2 N1 M0 | non-hispanic | White | F |
| 19 | 1A1266 | 3 | 70 | T3 N1 | non-hispanic | White | M |
| 20 | 1A1339 | 3.7 | 62 | T3 N1 | non-hispanic | White | M |
| 21 | 1A1425 | 1.8 | 62 | T3 N1 | non-hispanic | White | F |

**Supplementary Table S3: PDAC patient information for multiplex IF (Vectra®).**

| Number | Random Number | Tumor size (cm) | Age (y) | Stage | Race | Sex |
|---|---|---|---|---|---|---|
| 1 | 14977 | 1.7 | 77 | T1 N1 | Unknown | M |
| 2 | 14981 | 2.5 | 88 | T2 N2 | White | F |
| 3 | 14982 | 1.5 | 73 | T3a N0 | White | F |
| 4 | 14983 | 8 | 57 | T3 N0 | White | M |
| 5 | 14984 | 3 | 72 | T2 N2 | Unknown | M |
| 6 | 14985 | 7.2 | 65 | T3 N1 | White | M |
| 7 | 14987 | 3 | 84 | T2 N1 M1 | White | M |
| 8 | 14978 | 4.5 | 81 | T3 N1 | White | F |
| 9 | 14989 | 1.5 | 45 | T2 N2 | Other | F |
| 10 | 14990 | 0.1 | 48 | T1a N0 | White | M |
| 11 | 14991 | 3 | 71 | T3b N1 | Other | M |
| 12 | 14995 | 2.5 | 65 | T2 N0 | White | M |
| 13 | 14971 | 1.7 | 82 | T2 N1 | White | F |
| 14 | 14970 | 3 | 85 | T2 N1 | White | M |
| 15 | 14951 | 3.7 | 56 | T3 N1 | Unknown | M |
| 16 | 14946 | 1.6 | 71 | T1 N1 | White | F |
| 17 | 14979 | | | | | |

**Supplementary Table S4: Antibodies and fluorescent dyes for multiplex immunofluorescence (Vectra®).**

| Target | Company | Catalog Number | Dilution | Antigen retrieval | Fluorophore | Company |
|---|---|---|---|---|---|---|
| CD56 | Abcam | ab133345 | 1:100 | ER1 | Opal™ 570 | Akoya Biosciences Inc. |
| CD16 | Cell Signaling | 24326 | 1:300 | ER1 | Opal™ 480 | |
| Pan-cytokeratin | Abcam | ab185966 | 1:100 | ER1 | Opal™ 520 | |
| αSMA | Abcam | ab124964 | 1:500 | ER1 | Opal™ 690 | |
| CD335 | Thermofisher | PA5-79720 | 1:75 | ER1 | Opal™ 620 | |
| CD68 | Abcam | ab955 | 1:100 | ER2 | Opal™ 780 | |

**Supplementary Table S5: MMP genes primer sequences.**

| Gene | Forward Sequence (3'→5') | Reverse Sequence (3'→5') |
|---|---|---|
| *MMP1* | AAAATTACACGCCAGATTTGCC | GGTGTGACATTACTCCAGAGTTG |
| *MMP2* | TACAGGATCATTGGCTACACACC | GGTCACATCGCTCCAGACT |
| *MMP3* | AGTCTTCCAATCCTACTGTTGCT | TCCCCGTCACCTCCAATCC |
| *MMP7* | GAGTGAGCTACAGTGGGAACA | CTATGACGCGGGAGTTTAACAT |
| *MMP9* | TGCTCTGCCTATCCTCTGAGT | TCACATCCTTTTCGAGGTTGTAG |
| *MMP10* | TGCTCTGCCTATCCTCTGAGT | TCACATCCTTTTCGAGGTTGTAG |
| *MMP12* | CATGAACCGTGAGGATGTTGA | GCATGGGCTAGGATTCCACC |
| *MMP13* | ACTGAGAGGCTCCGAGAAATG | GAACCCCGCATCTTGGCTT |
| *MT1-MMP* | GGCTACAGCAATATGGCTACC | GATGGCCGCTGAGAGTGAC |
| *MT2-MMP* | AGGTCCATGCCGAGAACTG | GTCTCTTCGTCGAGCACACC |
| *MT3-MMP* | AGCACTGGAAGACGGTTGG | CTCCGTTCCGCAGACTGTA |
| *MT6-MMP* | GACTGGCTGACTCGCTATGG | TGATGGCATCGCGCAACTT |
